# Remodeling of sporozoite microtubules during *Plasmodium* liver stage development

**DOI:** 10.1101/2025.06.26.661843

**Authors:** Kodzo Atchou, Magali Roques, Ruth Rehmann, Reto Caldelari, Melanie Schmid, Simone Grossi, Bianca Manuela Berger, Torsten Ochsenreiter, Friedrich Frischknecht, Volker Heussler

**Affiliations:** Institute of Cell Biology, University of Bern, 3012 Bern, Switzerland; Graduate School for Cellular and Biomedical Sciences, University of Bern, 3012 Bern, Switzerland; Department of Parasitology, University of Heidelberg Medical School, Im Neuenheimer Feld 324, 69120, Heidelberg, Germany

## Abstract

Microtubules are essential components of the eukaryotic cytoskeleton, controlling cell shape, intracellular organization, and cell division. During the *Plasmodium* liver stage, parasites undergo exceptionally rapid nuclear replication, yet the organization and dynamics of their microtubule cytoskeleton remain poorly understood. Here, using expansion microscopy, live-cell imaging, and tubulin post-translational modification (PTM)-specific antibodies, we characterize microtubule organization throughout *Plasmodium berghei* liver stage development. We identified a prominent liver stage parasite microtubule bundle (LSPMB) that appears to originate from the sporozoite subpellicular microtubules. The LSPMB progressively shrinks during liver stage development and, in contrast to sporozoite subpellicular microtubules, becomes sensitive to microtubule depolymerization, revealing extensive remodeling of the parasite microtubule cytoskeleton. LSPMB filaments transiently associate with hemi-spindle poles during schizogony, suggesting that this pre-existing microtubule network may contribute to microtubule organization during nuclear replication. We further identify developmentally regulated changes in antibody signals associated with tubulin polyglutamylation and tyrosination and detect the microtubule plus-end-binding protein EB1 on both the LSPMB and hemi-spindles. Mutational analysis further revealed an important contribution of the α1-tubulin C-terminal region to parasite development: alteration of selected residues impaired liver stage growth, whereas deletion of the entire C-terminal tail severely compromised sporozoite motility and salivary gland colonization. Together, our findings reveal extensive remodeling of the *Plasmodium* microtubule cytoskeleton during the transition from the invasive sporozoite to the replicative liver stage and identify the α1-tubulin C-terminal region as an important regulator of parasite development and transmission.

## Introduction

*Plasmodium* parasites are responsible for malaria and cause approximately half a million deaths annually in sub-Saharan countries. Malaria continues to be a major worldwide health concern despite recent successful attempts to lower its prevalence (1–4). The infection starts when an infected female *Anopheles* mosquito, during a blood meal, injects salivary gland sporozoites into the skin of the vertebrate host. Some of the motile sporozoites escape the host immune defense and invade blood vessels. They are carried by the blood flow to the liver, where they invade and infect hepatocytes (5). Within the hepatocyte, one single parasite gives rise to thousands of daughter parasites called merozoites in 3 to 6 days depending on the *Plasmodium* species (6–8). Fully developed merozoites exit the liver cells in merosome vesicles through adjacent blood vessels (8–11). During liver stage development, the parasite relies on fast and massive asexual replication, which is essential for transmission and pathogenesis in the erythrocyte stage, representing key targets for malaria control strategies. Investigating the parasite microtubule dynamics during the rapid replication could provide important insight into parasite biology and help to develop strategies for halting the massive proliferation during the clinically asymptomatic liver stage.

Microtubules are cytoskeletal filaments of about 25 nm outer diameter, made up of usually 13 chains of protofilaments composed of globular α-tubulin and β-tubulin heterodimers. They are crucial components of the cytoskeleton and offer structural support to cells and function as tracks for the movement of organelles, vesicles, and other cellular components (12,13). Microtubules are involved in important cellular activities such as cell division and the production of cilia and flagella to allow cell motility (14,15). They exhibit inherent dynamics, alternating between polymerization (rapid growth) and spontaneous depolymerization (shrinkage) in a process termed dynamic instability (12,16,17). The dynamics of microtubules and their interactions with other cellular structures are regulated by specific microtubule-associated proteins (MAPs), which selectively bind to either the plus or minus ends of microtubules (18–20). The properties and functions of tubulin are largely encoded within their protein sequences, contributing to molecular diversity that influences key microtubule characteristics. This variation has been conceptualized as the "tubulin code,” a regulatory framework through which cells control microtubule diversity to achieve specific physiological functions (13). Tubulin is subject to at least seven identified post-translational modifications (PTMs). These include the acetylation of lysine 40 on α-tubulin, which faces the microtubule lumen, as well as glycylation, the addition of polyglycine side chains of varying lengths. Other modifications are phosphorylation, palmitoylation, and the cycle of tyrosination/detyrosination, the latter involving the addition or removal of a C-terminal tyrosine and often linked to the regulation of dynamic versus stable microtubules (21). Stable microtubules represent a specialized subset with increased resistance to depolymerization, even under destabilizing conditions. Additionally, glutamylation (or polyglutamylation) entails the addition of multiple glutamate side chains to glutamate residues on both α- and β-tubulin, primarily within their C-terminal tails, and has been associated with stable microtubules (22–25). This process is catalyzed by enzymes known as polyglutamylases (26,27). These PTM-related enzymes are highly conserved and allow cells to fine-tune microtubule properties, dynamics, and interactions with MAPs (28,29). Nevertheless, the precise mechanisms cells use to establish and interpret this code, particularly in *Plasmodium*, are not yet fully understood. Although highly conserved across evolution, tubulin exhibits structural and functional variations between species, cell types, and even within single cells. In yeast, two distinct genes encode α-tubulin, while a single gene encodes β-tubulin. In contrast, the human genome contains nine genes encoding α-tubulin and an equal number encoding β-tubulin, reflecting a greater level of complexity and functional specialization in higher eukaryotes (13,30,31). Similar to yeast, the *Plasmodium* genome encodes two α-tubulins (α1-tubulin and α2-tubulin) and one β-tubulin (32–36). The α1-tubulin gene encodes a protein of 453 amino acids, while α2-tubulin encodes 450 amino acids (37). The two isoforms share 94% amino acid identity. However, α2-tubulin lacks a terminal tyrosine residue, which is present in α1-tubulin. This difference suggests potential variation in post-translational modifications and functional roles (33,38). In *P. berghei,* the α1-tubulin gene is expressed across most stages of the *Plasmodium* life cycle, whereas α2-tubulin expression appears restricted to male gametocytes, gametes, and newly formed zygotes, indicating distinct functional roles for the two isoforms (29,39). Deletion of α1-tubulin blocks sporozoite formation in oocysts and lowering α1-tubulin expression leads to less infectious sporozoites (39). There is a significant degree of amino acid conservation between human and *Plasmodium* tubulins, with approximately 83.7% identity for α-tubulin and 88.5% for β-tubulin. However, *Plasmodium* tubulin shows greater similarity to plant tubulin than to mammalian tubulin. This raises the potential for developing compounds that selectively target parasite microtubules while sparing the host cell cytoskeleton, offering a promising avenue for therapeutic intervention (32,34–36,40).

Recent studies on *Plasmodium falciparum* blood stage schizonts have revealed that hemi-spindles lack polyglutamylation, whereas subpellicular microtubules do exhibit this post-translational modification (41–43). Similarly, SPMTs appear to be stabilized by microtubule inner proteins, while spindle microtubules do not show these luminal proteins (44). These differences may reflect functional specialization of microtubule subpopulations in different cellular structures and developmental stages.

The liver stage of *Plasmodium* exhibits one of the fastest rates of replication known among eukaryotic cells, a process likely dependent on microtubule dynamics. However, to date, no studies have investigated microtubule dynamics during the liver stage. This study was therefore designed to explore the dynamics of parasite tubulin during the asexual liver stage, using molecular biology approaches and recently developed expansion microscopy protocols (43,45–47).

## Results

### Expansion microscopy reveals the presence of the sporozoite subpellicular microtubules during the liver stage of infection

To investigate microtubule organization during the *Plasmodium* parasite liver stage, HeLa cells were infected with sporozoites and fixed at different time points using cold methanol. We then performed IFA to stain host and parasite microtubules using an *α*-tubulin antibody (Figure 1A). The IFA revealed a marked accumulation of microtubules in proximity to the parasitophorous vacuole membrane (PVM) (stained with anti-UIS4) from early infection (24 hours post-infection (hpi)) to schizogony (30 to 56 hpi) (Figure 1A insets). To examine these structures in greater detail, we applied the recently developed expansion microscopy protocols (PS-ExM and U-ExM) (45–47). Using PS-ExM, both host and parasite microtubules were successfully expanded five-fold while preserving their structure (45). In contrast, the U-ExM protocol (46,47) did not preserve host cell microtubules, likely due to denaturation or epitope loss during processing. However, this protocol successfully retained parasite microtubules. This selective preservation helped distinguish parasite microtubules from those of the host, allowing us to focus specifically on the parasite’s cytoskeleton. By expanding the cells, U-ExM enabled the visualization of a prominent tubulin structure within the parasite, detectable from 6 to 48 hpi (Figure 1B). Based on observations of sporozoites prior to liver infection, we hypothesize that this structure originates from the sporozoite subpellicular microtubules (SSPMTs) (Figure 1C top panel). We refer to this structure as the liver stage parasite microtubule bundle (LSPMB). Notably, the LSPMB progressively shrank over time and was no longer detectable by the end of schizogony (56 hpi) (Figure 1A, B). Using the 3D and 4D imaging software IMARIS, we quantified the LSPMB volume at various stages of liver infection. The results confirmed a consistent decrease in LSPMB size between 6 and 56 hpi (Figure 1D).

To verify that the LSPMB is not an artifact of the IFA and PS-ExM, we performed live-cell imaging of infected HeLa cells. Cells were incubated with the tubulin tracker SPY650 for one hour prior to imaging. The LSPMB was clearly visible in sporozoites before invasion (Figure 1–video 1) and persisted until late liver stage development, gradually shrinking during schizogony (Figure 1E; Figure 1–video 2, 3, and 4, ∼36 hpi to 48 hpi). These live imaging results corroborate our findings from IFA and expansion microscopy, validating the presence of the LSPMB during liver stage development. The pronounced changes in LSPMB morphology observed by live imaging prompted us to investigate more closely whether these structures retain the remarkable stability of sporozoite subpellicular microtubules or become dynamic during liver stage development.

**Figure 1.**
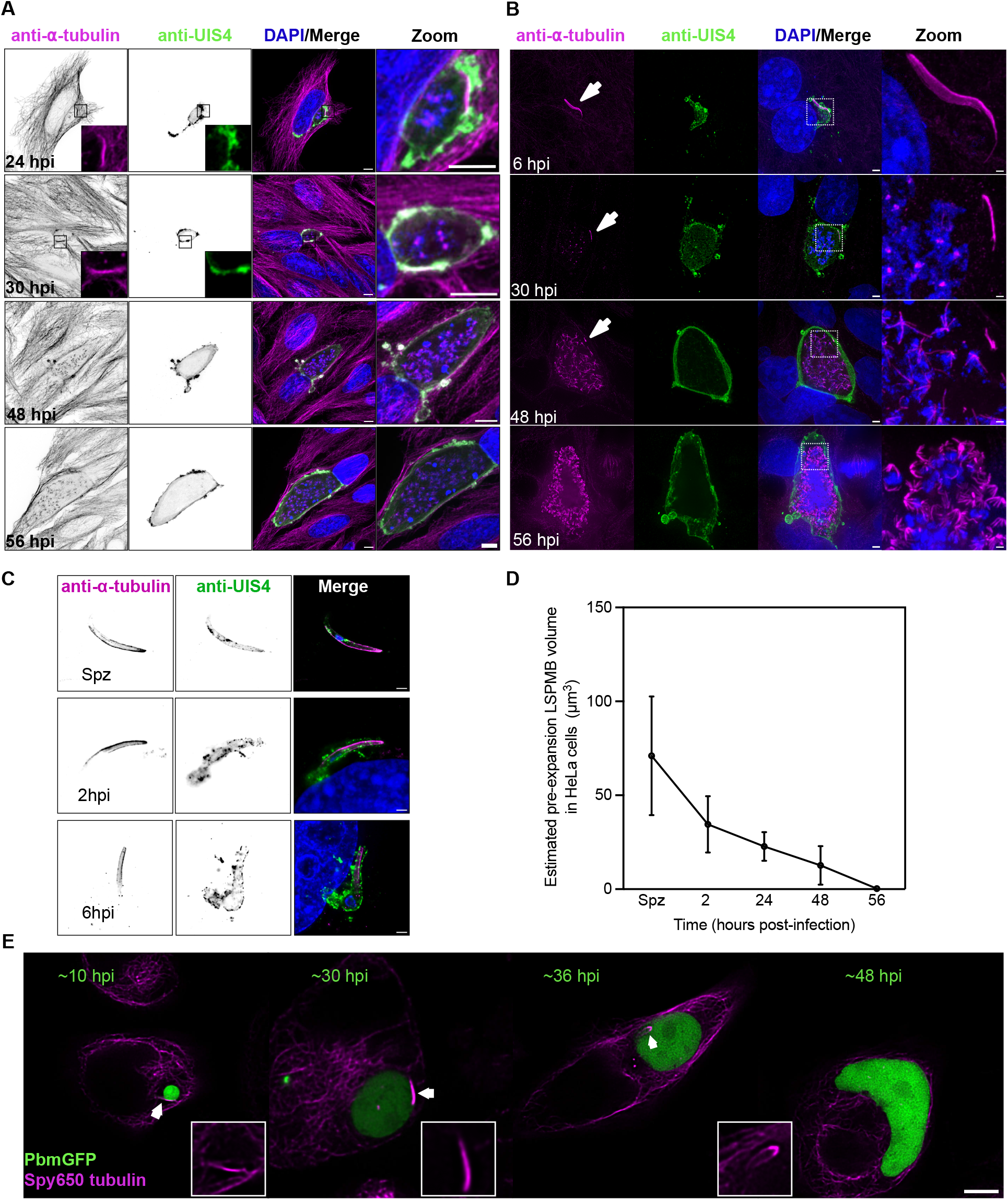
Immunofluorescence and expansion microscopy reveal remodeling of sporozoite microtubules during liver stage development. **(A, B)** Maximum-intensity projections of confocal z-stacks of *Plasmodium berghei*-infected HeLa cells. Host and parasite microtubules were labelled with an anti-α-tubulin antibody (magenta), nuclei were stained with DAPI (blue), and the PVM was labelled with an anti-UIS4 antibody (green). **(A)** HeLa cells were infected with salivary gland sporozoites and methanol-fixed at 24, 30, 48 and 56 hpi. Insets at 24 and 30 hpi highlight microtubule accumulation around the PVM. The zoomed panels show magnified views of the parasites. Scale bars, 5 µm; zoomed panels, 2 µm. **(B)** Infected HeLa cells were fixed at 6, 30, 48 and 56 hpi and expanded using the U-ExM protocol. Insets and white arrows indicate the liver stage parasite microtubule bundle (LSPMB). The zoomed panels show magnified views of the parasite microtubules. Scale bars, 10 µm; zoomed panels, 2 µm. **(C)** Maximum-intensity projections of expanded salivary gland sporozoites and intracellular parasites at 2 and 6 hpi, illustrating the relationship between the sporozoite subpellicular microtubules (SSPMTs) and the early LSPMB. Microtubules were labelled with an anti-α-tubulin antibody, and the PVM was labelled with an anti-UIS4 antibody. Spz, salivary gland sporozoite. Scale bars, 10 µm. **(D)** Quantification of the segmented anti-α-tubulin-positive SSPMT/LSPMB signal volume from expanded confocal z-stacks using three-dimensional surface reconstruction in Imaris. Volumes were divided by the cube of the linear expansion factor (5³ = 125) to estimate the corresponding pre-expansion volumes. Mean volumes were 71 µm³ in salivary gland sporozoites and 34.48, 22.76, 12.69 and 0.32 µm³ at 2, 24, 48 and 56 hpi, respectively. Data are presented as mean ± SD; n = 15 parasites per time point from three independent experiments. **(E)** Representative live-cell confocal images of infected HeLa cells incubated with SPY650-tubulin. Images were acquired at approximately 10, 30, 36 and 48 hpi. Host and parasite microtubules are shown in magenta, and parasites express cytosolic GFP (green). White arrows indicate the LSPMB, and boxed regions indicate the corresponding magnified views. Scale bars, 5 µm. All experiments were performed in three independent biological replicates.

**Figure 1–video 1**: Time-lapse live imaging of *P. berghei* salivary gland sporozoites expressing GFP cytosolically. Sporozoites were isolated and incubated with tubulin tracker SPY650. Microtubules are shown in magenta; sporozoites are shown in green. Scale bar: 5 µm.

**Figure 1–video 2, 3**, and **4**: Time-lapse live imaging of HeLa cells infected with *P. berghei* liver stage parasites expressing cytosolic GFP. Infected cells were incubated with tubulin tracker SPY650 for one hour prior to imaging. The movies show the development and dynamics of the LSPMB during liver stage infection. Microtubules are shown in magenta; parasites are shown in green. **Video 2**: imaging from 10 hpi to 25 hpi. **Video 3:** imaging from 30 hpi to 35 hpi, and **Video 4:** imaging from 37 hpi to 49 hpi. Scale bar: 5 µm.

### The liver stage parasite subpellicular microtubules are dynamic

The SSPMTs have previously been described as highly stable structures, enabling sporozoite formation and invasion (39,44,48). These microtubules, similar to the corresponding microtubules in *T. gondii*, are resistant to conventional depolymerizing agents (48). To investigate whether the LSPMB retains this stability during liver stage development, we treated infected HeLa cells with the microtubule-depolymerizing agents Amiprophos-methyl (AMP) and the herbicide oryzalin. These compounds are known to selectively target dynamic *Plasmodium* microtubules affecting mammalian cells only in high concentrations (for AMP 50 µM) (32,34,36,40,49) (Figure 2 and Figure 2–figure supplement 1A and B). We first validated drug specificity and cytotoxicity by testing various concentrations on HeLa cells over one week. Oryzalin did not significantly affect HeLa cell numbers at the tested concentrations, whereas 50 µM AMP reduced cell numbers relative to the DMSO control (Figure 2–figure supplement 1A and B) or disrupted host microtubules (Figure 2–figure supplement 1D and E). However, oryzalin showed a stronger inhibitory effect on parasite growth compared to AMP (Figure 2–figure supplement 1C) and was therefore selected for further experiments (Figure 2A). Mosquito salivary glands were dissected and sporozoites collected in medium containing 10 µM or 20 µM oryzalin and then used to infect HeLa cells (Figure 2A and B), which were incubated with the same drug concentrations. At 48 hpi, the cells were fixed and analyzed by IFA and PS-ExM. To test reversibility, oryzalin was washed out at 24 hpi prior to the onset of schizogony (Figure 2B). At 10 µM, oryzalin significantly impaired nuclear replication, spindle formation, and LSPMB volume when present throughout the culture period (Figure 2C, D and E). Upon drug removal, partial recovery of hemi-spindles and nuclear replication was observed, suggesting that the effect of 10 µM oryzalin is reversible at this stage (Figure 2C, D, and E). However, at 20 µM, neither hemi-spindles nor the LSPMB showed detectable recovery after drug removal, although general parasite damage at this concentration may have contributed to the lack of recovery (Figure 2A and B). To investigate whether the microtubule depolymerization by oryzalin affects parasite growth and survival during the liver stage, we measured parasite size within infected HeLa cells at 6, 24 and 48 hpi. Treatment with 10 µM oryzalin led to a significant reduction in parasite size and survival (Figure 2 F, G, H, and I). We further examined merozoite formation by staining for MSP1 at 56 hpi. Following continuous treatment with 10 µM oryzalin, MSP1 signal was not detected, and neither hemi-spindle nor LSPMB structures were restored, consistent with severely impaired merozoite development.

This suggests that intact microtubules are essential for DNA replication and merozoite formation (Figure 2J). Interestingly, at lower concentrations of 0.5 µM, oryzalin treatment led to an unexpected elongation of the LSPMB and hemi-spindles up to 3.4 times longer than controls, suggesting that microtubule turnover is tightly regulated (Figure 2–figure supplement 1F, G and H). In conclusion, their sensitivity to oryzalin is consistent with increased susceptibility to microtubule disruption during liver stage development. This dynamic behavior was further supported by live-cell imaging using the SPY650 tubulin tracker (Figure 2– video 1, 2, and 3). Given the observed microtubule remodeling in response to oryzalin, we next sought to determine whether known microtubule-associated proteins, specifically those regulating plus-end dynamics, are expressed during the liver stage and might contribute to the observed plasticity. Only a limited number of motor proteins and microtubule-associated proteins, such as centrin and the kinesin protein family, have been identified during the *Plasmodium* liver stage (8,50–52). The recently characterized EB1 protein, known to regulate microtubule plus-end dynamics and length, has so far been reported as being expressed primarily during the sexual blood stage (53–55). However, these studies did not assess EB1 expression during the mosquito and liver stage. Given our observations of dynamic microtubule behavior and excessive polymerization following oryzalin treatment, we explored whether EB1 might be expressed during the liver stage. To this end, we generated a parasite line expressing endogenous EB1 fused to eGFP via double homologous recombination (Figure 2–figure supplement 2C and D; Figure 2– figure supplement 2–Source Data 1). As expected, EB1-eGFP was detected in sexual blood stages (Figure 2–figure supplement 2E and F) consistent with previous reports (53,54). Interestingly, we also detected EB1–eGFP in oocysts at 12 days after the infectious blood meal and in salivary gland sporozoites (Figure 2–figure supplement 2G, H). During liver stage development, EB1–eGFP was associated with the LSPMB and hemi-spindles (Figure 2–figure supplement 2I). This localization is consistent with a potential role in parasite microtubule organization, although the function of EB1 during these stages remains to be established.

**Figure 2.**
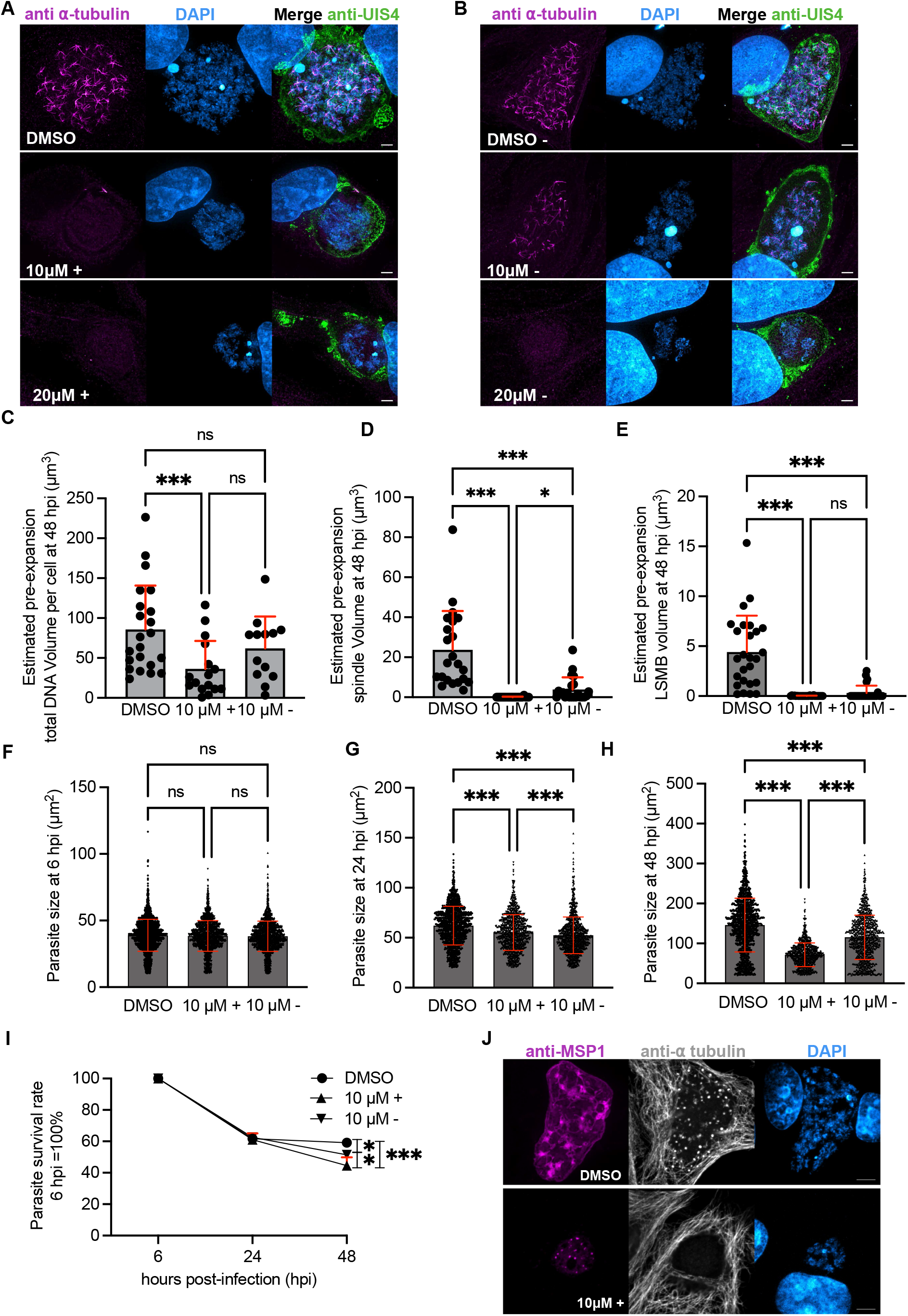
Oryzalin disrupts microtubule organization and impairs *Plasmodium* liver stage development. **(A, B)** Maximum-intensity projections of confocal z-stacks of expanded *Plasmodium berghei*-infected HeLa cells. Salivary gland sporozoites were used for infection, and cells were cultured in the presence of DMSO or 10 or 20 µM oryzalin. Cells were methanol-fixed and expanded at 48 hpi. Parasite and host microtubules were labelled with an anti-α-tubulin antibody (magenta), the PVM was labelled with an anti-UIS4 antibody (green), and nuclei were stained with DAPI (blue). **(A)** DMSO control and continuous treatment with 10 or 20 µM oryzalin until fixation at 48 hpi. **(B)** Corresponding washout conditions in which oryzalin was removed at 24 hpi and cells were cultured in drug-free medium until fixation at 48 hpi. Scale bars, 10 µm. In the figure, “+” indicates continuous treatment and “−” indicates washout at 24 hpi. **(C)** Estimated pre-expansion volume of the segmented DAPI-positive parasite nuclear signal at 48 hpi. Mean volumes were 85.7 µm³ for DMSO-treated parasites, 36.3 µm³ following continuous treatment with 10 µM oryzalin, and 62.0 µm³ following washout at 24 hpi. n = 22 parasites per condition. **(D)** Estimated pre-expansion total hemi-spindle signal volume at 48 hpi. Mean volumes were 23.7 µm³ for DMSO-treated parasites and 3.9 µm³ following washout. No hemi-spindle signal satisfying the segmentation criteria was detected following continuous treatment with 10 µM oryzalin. n = 22 parasites per condition. **(E)** Estimated pre-expansion LSPMB signal volume at 48 hpi. Mean volumes were 4.4 µm³ for DMSO-treated parasites, 0.01 µm³ following continuous treatment with 10 µM oryzalin, and 0.3 µm³ following washout. n = 26 parasites per condition. For **(C–E)**, volumes measured from expanded samples were divided by the cube of the linear expansion factor (5³ = 125). **(F–H)** Parasite areas at 6 **(F)**, 24 **(G)** and 48 hpi **(H)** under the indicated treatment conditions. Parasite sizes were comparable at 6 hpi. Continuous oryzalin treatment reduced parasite growth at 24 and 48 hpi, whereas washout at 24 hpi resulted in partial recovery. Each dot represents one parasite. **(I)** Parasite survival between 6 and 48 hpi, normalized to the corresponding parasite number at 6 hpi. Continuous treatment with 10 µM oryzalin reduced parasite survival at 48 hpi relative to the DMSO control, whereas washout at 24 hpi resulted in partial recovery. Data represent mean ± SD from three independent experiments. **(J)** Representative maximum-intensity projections of confocal z-stacks of infected HeLa cells at 56 hpi following continuous treatment with DMSO or 10 µM oryzalin. Cells were methanol-fixed and labelled with antibodies against MSP1 (magenta) and α-tubulin (grey); nuclei were stained with DAPI (blue). Scale bars, 5 µm. In **(C–H)**, each dot represents one parasite, and red summary lines indicate mean ± SD. All experiments were performed in three independent biological replicates. Statistical analysis was performed using one-way ANOVA followed by (Tukey’s multiple-comparisons test for all pairwise comparisons/Dunnett’s multiple-comparisons test for comparisons with DMSO). ns, not significant; *p ≤ 0.05; **p ≤ 0.01; ***p ≤ 0.001.

**Figure 2–video 1**, **2**, and **3** Time-lapse live imaging of HeLa cells infected with *P. berghei* liver stage parasites expressing cytosolic GFP. Infected cells were incubated with tubulin tracker SPY650 for one hour prior to imaging. The movies highlight the high dynamicity of the LSPMB during liver stage development, starting from early schizogony (∼30 hpi). Microtubules are shown in magenta; parasites are shown in green. Scale bar: 5 µm.

### Developmental changes in tubulin PTM-associated antibody signals and LSPMB organization

Microtubule functions are often regulated by different PTMs, which influence their stability, interactions, and dynamics. To characterize developmental changes in the LSPMB, we used immunofluorescence, expansion microscopy and live microscopy in infected HeLa cells (Figure 3 and Figure 2–video 1, 2 and 3). To investigate whether the LSPMB is present under more physiological conditions and not only in model HeLa cells, we infected isolated primary mouse hepatocytes with *Plasmodium* sporozoites and fixed the cells at different time points (6, 24, 36, 48, 56 and 60 hpi). We also analyzed sporozoites isolated from mosquito salivary glands using antibodies specific for polyglutamylated α-tubulin (anti-IN105, herein referred to as anti-PolyE) (56) (Figure 3A) and tyrosinated α-tubulin (anti-T9028, referred to as anti-Ty, and anti-YL1/2) (Figure 3B and Figure 3–figure supplement 1). Using these markers, we detected both anti-PolyE and anti-Ty signals along sporozoite SSPMTs and the early LSPMB. Quantitative colocalization analysis showed a strong spatial overlap between the anti-PolyE and anti-α-tubulin signals defining the SSPMTs and LSPMB in sporozoites and liver stage parasites, respectively (Figure 2–figure supplement 2A).

Interestingly, following hepatocyte invasion, the anti-Ty signal on the LSPMB became undetectable, whereas anti-PolyE signals remained detectable. The loss of anti-Ty signal persisted throughout liver stage development, including early and late schizogony (24–56 hpi). During this period, anti-PolyE signals were detected on the LSPMB and at hemi-spindle poles, but not along the polymerizing hemi-spindles themselves, which eventually disappeared along with the LSPMB (Figure 3A and C). Given previous findings observing polyglutamylation in hemi-spindle pole formation in erythrocytic merozoites (28), we investigated whether there is a structural relationship between the LSPMB and the hemi-spindle poles. Using PS-ExM and the tubulin tracker SPY650, we observed LSPMB filaments closely opposed to hemi-spindle poles. Live-cell imaging revealed transient spatial associations between these structures, particularly during LSPMB shrinkage (Figure 3F; Figure 3–video 1, 2 and 3, Figure 3–video supplement 1). These observations raise the possibility that LSPMB-derived tubulin may contribute to hemi-spindle organization during schizogony. We also stained for γ-tubulin and observed its localization at the hemi-spindle poles in both the parasite and host cell (Figure 2–figure supplement 2B), supporting the idea that *Plasmodium* may utilize γ-tubulin for nuclear division and microtubule assembly, as reported in other systems (57–60).

To further define the spatial localization of the LSPMB, we performed PS-ExM using a previously generated *P. berghei* line expressing the parasite plasma membrane protein PMP1 fused to superfolder GFP (sfGFP) (61). Fluorescence-intensity profiling of the expanded samples positioned the LSPMB signal close to the parasite periphery, between or adjacent to the signals marking the PMP1-labelled parasite plasma membrane and the UIS4-labelled PVM (Figure 3D and E). Although PS-ExM provides substantially improved spatial resolution, it does not resolve the precise membrane topology of the LSPMB; this will require higher-resolution ultrastructural approaches. At later stages, particularly during cytomere formation, anti-Ty signals reappeared near parasite nuclei and subsequently appeared as punctate structures within forming merozoites (Figure 3B).

Tyrosination has previously been associated with dynamic microtubules (39,44,62,63). The absence of detectable anti-Ty signal during early intracellular liver stage development coincided with the transformation of the slender, motile sporozoite into a rounded, replicating liver stage parasite (Figure 1 and 3). The developmental loss of detectable anti-Ty signal may accompany LSPMB remodeling during the liver stage, potentially making tubulin available for hemi-spindle formation during schizogony.

The reappearance of punctate anti-Ty signals during merozoite formation may accompany microtubule reorganization at this developmental stage, although the molecular identity and functional significance of these signals remain unresolved.

**Figure 3.**
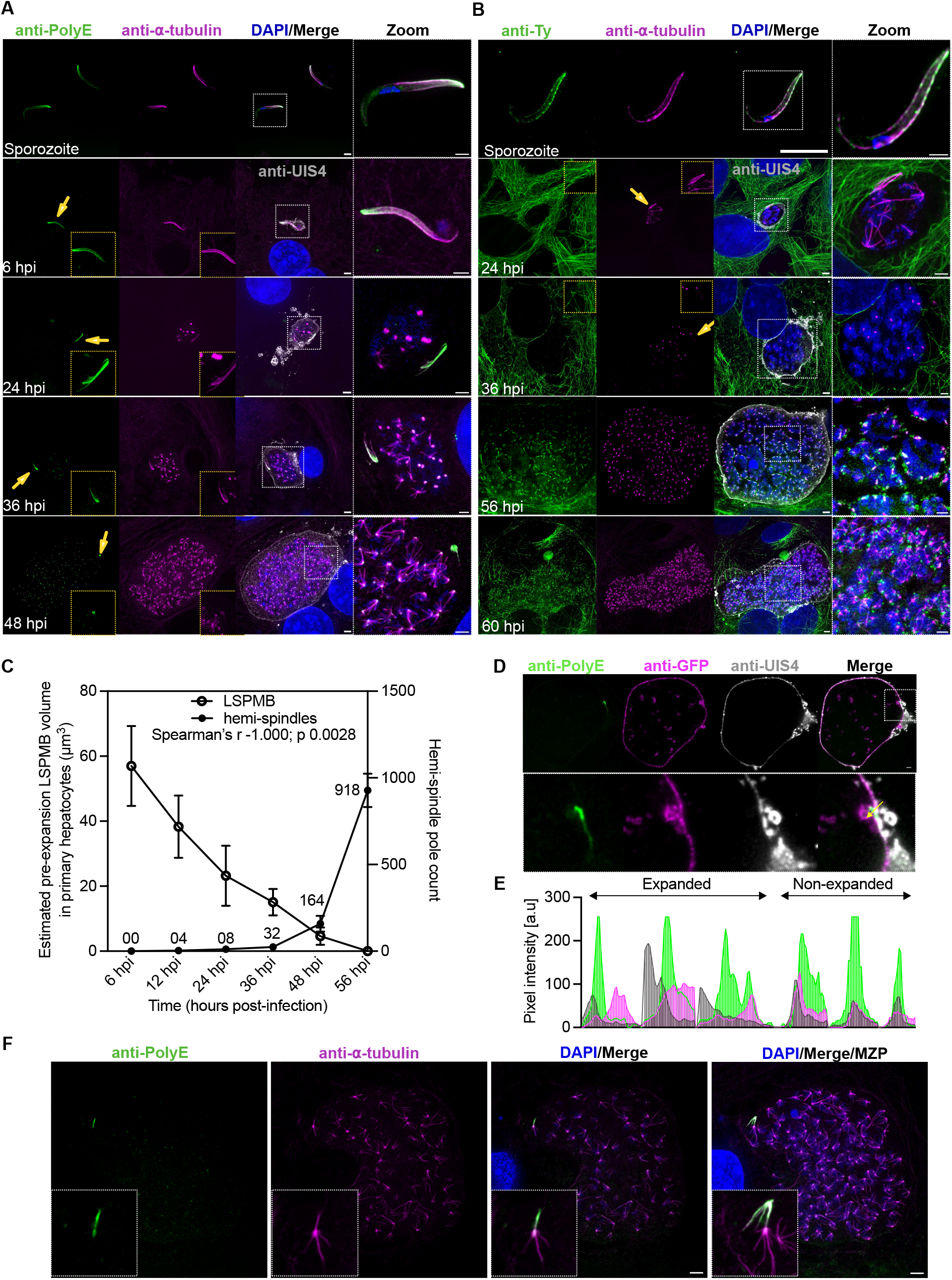
Developmental changes in tubulin PTM-associated antibody signals and LSPMB organization. **(A, B)** Maximum-intensity projections of confocal z-stacks acquired by PS-ExM from salivary gland sporozoites and infected primary hepatocytes at the indicated stages of liver stage development. Primary hepatocytes were isolated from mice and infected with *Plasmodium berghei* sporozoites. Samples were fixed and labelled with **(A)** an anti-PolyE antibody (IN105) or **(B)** an anti-Ty antibody (T9028), together with an anti-α-tubulin antibody. Nuclei were stained with DAPI (blue), and the PVM was labelled with an anti-UIS4 antibody (grey). Yellow arrows and insets highlight remodeling of the LSPMB and the appearance of anti-PolyE antibody signals at hemi-spindle poles. The zoomed panels show magnified views of the LSPMB and other parasite microtubule structures. Scale bars, 10 µm; boxed magnified regions, 5 µm. **(C)** Estimated pre-expansion LSPMB signal volume and mean hemi-spindle pole number per parasite between 6 and 56 hpi, quantified by three-dimensional analysis in Imaris. Volumes measured from expanded samples were divided by the cube of the linear expansion factor (5³ = 125). The mean LSPMB signal volume decreased from 56.97 µm³ at 6 hpi until no LSPMB signal satisfying the segmentation criteria was detected at 56 hpi. In parallel, the mean number of hemi-spindle poles increased from 0 at 6 hpi to 918 at 56 hpi. The displayed numbers indicate the mean hemi-spindle pole count at each time point. Data are presented as mean ± SD; n = 15 parasites per time point from three independent experiments. Across the six time-point means, LSPMB signal volume and hemi-spindle pole number showed a perfect negative rank correlation (Spearman’s ρ = −1.00, *P* = 0.0028; n = 6 time points). **(D)** Representative single-plane confocal image of an expanded *P. berghei*-infected HeLa cell at 36 hpi expressing the parasite plasma membrane marker PMP1::GFP. Cells were PFA-fixed and labelled with anti-PolyE (green), anti-GFP (magenta) and anti-UIS4 (grey) antibodies. The yellow arrow indicates the direction of the line scan from outside the PVM towards the parasite interior. The inset shows a magnified view of the subcellular position of the LSPMB-associated anti-PolyE signal. Scale bar, 10 µm. **(E)** Pixel-intensity profiles of the anti-UIS4-labelled PVM (grey), PMP1::GFP-labelled parasite plasma membrane (magenta), and anti-PolyE signal associated with the LSPMB (green) in expanded and non-expanded samples. In non-expanded samples, the three signals could not be spatially resolved. Following PS-ExM, the anti-PolyE signal peak was spatially separated from the two membrane-marker signals and was detected within the interval bounded by them. For visualization, line scans from ten cells were concatenated for each condition. Data were obtained from three independent experiments. **(F)** Representative single-plane confocal image of an infected primary hepatocyte fixed with cold methanol at 48 hpi. Samples were labelled with anti-PolyE (green) and anti-α-tubulin (magenta) antibodies, and nuclei were stained with DAPI (blue). The region of interest shows an LSPMB filament closely apposed to a parasite hemi-spindle pole. MZP, maximum-intensity z-projection. Scale bar, 10 µm. Images are representative of three independent experiments.

**Figure 3–video 1**, **2**, and **3** Time-lapse live imaging of HeLa cells infected with *P. berghei* liver stage parasites expressing cytosolic GFP. Infected cells were incubated with tubulin tracker SPY650 for one hour prior to imaging. The movies reveal the spatial association between the LSPMB and the parasite’s hemi-spindles and spindle poles during liver stage development. Microtubules are shown in magenta; parasites are shown in green. **Video 1**: LSPMB (white arrow) localizes at the periphery of the parasite and is closely associated with hemi-spindles. **Video 2:** LSPMB filaments (white arrows) are transiently found in close spatial association with parasite hemi-spindle poles. **Video 3:** shrinkage of the LSPMB during liver stage development (ROI marked by a white arrow). Scale bar: 5 µm

### The α1-tubulin C-terminal region contributes to sporozoite development

To investigate the functional contribution of the α1-tubulin C-terminal region during liver stage development, we generated *P. berghei* mutant lines carrying targeted alterations in residues associated with the anti-PolyE and anti-Ty antibody epitopes (Figure 4A and Figure 4–figure supplement 1A, B). We generated three mutant lines: PsΔ, in which the predicted polyglutamylation-associated residue E445 was replaced with alanine; TyΔ, in which the C-terminal YEADY motif was deleted from position 449; and CtΔ, in which the entire C-terminal tail sequence EGEDEGYEADY was deleted (Figure 4A and Figure 4–figure supplement 1A, B).

**Figure 4:**
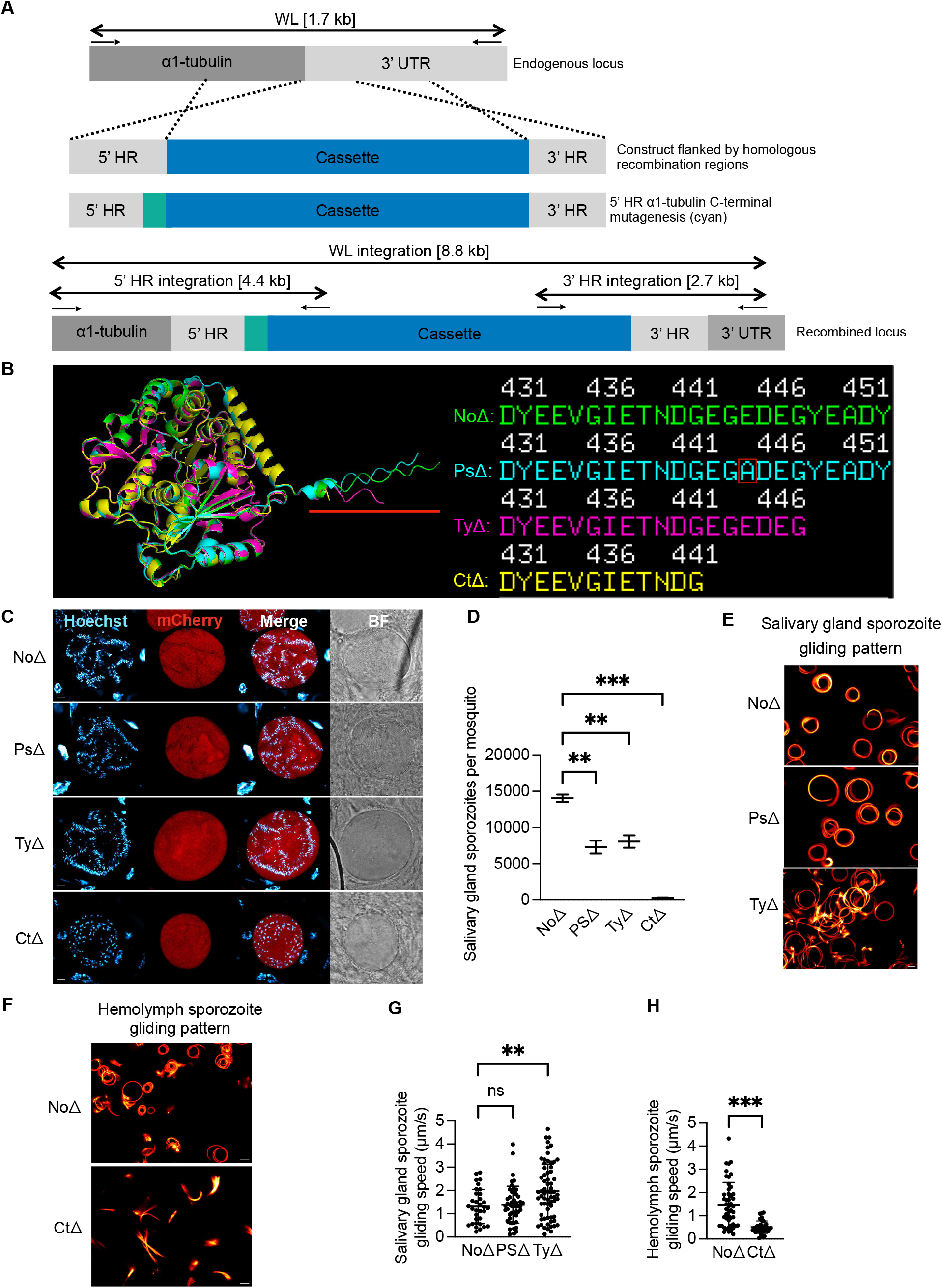
The α1-tubulin C-terminal region contributes to *Plasmodium* mosquito stage development. **(A)** Schematic of the double homologous recombination construct targeting the *Plasmodium berghei* α1-tubulin gene (*PBANKA_0417700*) for C-terminal mutagenesis. The cassette (blue) comprises the 3′ UTR of *P. berghei* dihydrofolate reductase (*PbDHFR*), the elongation factor 1α (*ef1α*) promoter, human dihydrofolate reductase (*hdhfr*), yeast cytosine deaminase/uracil phosphoribosyltransferase (*yfcu*), the 3′ UTR of *PbDHFR*, the Hsp70 promoter, mCherry, and the Hsp70 3′ UTR. The cassette was flanked by a 5′ homologous region (5′ HR) containing the α1-tubulin coding region, including the stop codon and the introduced C-terminal mutations (cyan), and a 3′ homologous region (3′ HR) comprising part of the *PBANKA_0417700* 3′ UTR. The construct was transfected into PbWT (*P. berghei* ANKA cl15cy1). Black arrows indicate PCR primer-binding sites and the expected product sizes. Primer sequences are listed in Tables 2 and 3. WL, whole locus. **(B)** Structural comparison of the α1-tubulin mutant proteins based on AlphaFold 3 predictions and alignment using PyMOL version 2.6.0. Green, control with an unaltered α1-tubulin C-terminal sequence (NoΔ); cyan, PsΔ; magenta, TyΔ; yellow, CtΔ. The predicted global folds of the four proteins were similar. The corresponding C-terminal amino acid sequences are shown on the right. **(C)** Representative confocal images of live oocysts at 11 days post-infection (dpi). Nuclei were stained with Hoechst 33342 (cyan), and cytosolic mCherry (red) was expressed under the Hsp70 promoter. BF, bright field. Scale bar, 5 µm. **(D)** Salivary gland sporozoite numbers per mosquito at 18 dpi. Mean numbers were 14,033 for NoΔ, 7,300 for PsΔ, 8,066 for TyΔ, and 200 for CtΔ. Data are presented as mean ± SD from three independent mosquito infections, with 20 mosquitoes analyzed per infection. Statistical analysis was performed using one-way ANOVA followed by Dunnett’s multiple-comparisons test. **p ≤ 0.01; ***p ≤ 0.001. **(E, F)** Representative temporally color-coded maximum-intensity projections of time-lapse motility assays generated using Fiji. Salivary gland sporozoites of the NoΔ, PsΔ, and TyΔ lines showed circular gliding patterns **(E)**. Because the very low number of CtΔ salivary gland sporozoites precluded a reliable motility analysis, CtΔ motility was examined using hemolymph sporozoites **(F)**. Under these conditions, CtΔ hemolymph sporozoites showed strongly impaired gliding and rarely formed the circular trails observed for NoΔ controls. Scale bars, 5 µm. **(G)** Gliding speeds of NoΔ, PsΔ, and TyΔ salivary gland sporozoites. Mean velocities were 1.30, 1.38, and 1.90 µm/s, respectively. Each dot represents one sporozoite, and data were obtained from three independent experiments. Statistical analysis was performed using one-way ANOVA followed by Dunnett’s multiple-comparisons test. ns, not significant; **p ≤ 0.01. **(H)** Gliding speeds of NoΔ and CtΔ hemolymph sporozoites. Mean velocities were 1.40 and 0.50 µm/s, respectively. Each dot represents one sporozoite, and data were obtained from three independent experiments. Statistical analysis was performed using a two-tailed unpaired *t*-test. ***p ≤ 0.001.

AlphaFold predictions suggested that the mutations were unlikely to cause major changes in overall tubulin structure (Figure 4B). The mutants were generated via double homologous recombination and verified by integration PCR (Figure 4–figure supplement 1C, Figure 4-figure supplement 1–Source Data 1) and sequencing of the mutated α-tubulin gene (Figure 4–figure supplement 1B). The development of mutant parasites was investigated during the mosquito and liver stages. Significant defects emerged during mosquito stage development. All three mutant lines formed oocysts and produced midgut sporozoites, although CtΔ showed altered oocyst development and a markedly reduced number of midgut sporozoites (Figure 4C; Figure 5–figure supplement 1A and B). PsΔ and TyΔ produced motile salivary gland sporozoites, albeit in reduced numbers, whereas only very few CtΔ sporozoites reached the salivary glands (Figure 4D–H; Figure 4–video supplement 1). Quantification at 18 days after the infectious blood meal showed that PsΔ and TyΔ produced approximately half as many salivary gland sporozoites as the NoΔ control, while CtΔ produced approximately 200 sporozoites per mosquito compared with approximately 14,000 for NoΔ (Figure 4D). We next examined sporozoite motility, which typically consists of continuous circular gliding (64–69). PsΔ and TyΔ salivary gland sporozoites were compared with stage-matched NoΔ controls. PsΔ sporozoites showed a gliding velocity similar to that of NoΔ, whereas TyΔ sporozoites displayed an increased velocity (1.9 versus 1.3 µm/s; Figure 4G), indicating that deletion of the C-terminal YEADY motif alters sporozoite motility.

To assess whether these motility phenotypes influenced infectivity, we performed typical infection assays in mice with PsΔ and TyΔ. Sporozoites of both mutant lines were able to infect mice similarly to control parasites (Figure 5–figure supplement 1C). Since CtΔ mutant parasites produced very few salivary gland sporozoites, we analyzed hemolymph sporozoites for this line compared to the controls. These showed severely impaired motility, lacking the typical gliding pattern and exhibiting significantly reduced speed (Figure 4F and H). Given that PsΔ and TyΔ mutants produced motile salivary gland sporozoites, we next infected HeLa cells with PsΔ and TyΔ mutant sporozoites and CtΔ mutant hemolymph sporozoites to assess their liver stage development and identify any downstream phenotypic effects.

### The *Plasmodium α*-tubulin C-terminal mutants undergo liver stage development

Using the anti-PolyE antibody, we detected signals associated with SSPMTs in PsΔ sporozoites and with the LSPMB during liver stage development (Figure 5A–C). Quantification at 36 hpi showed significantly reduced anti-PolyE fluorescence in PsΔ parasites compared with NoΔ controls (Figure 5G).

**Figure 5.**
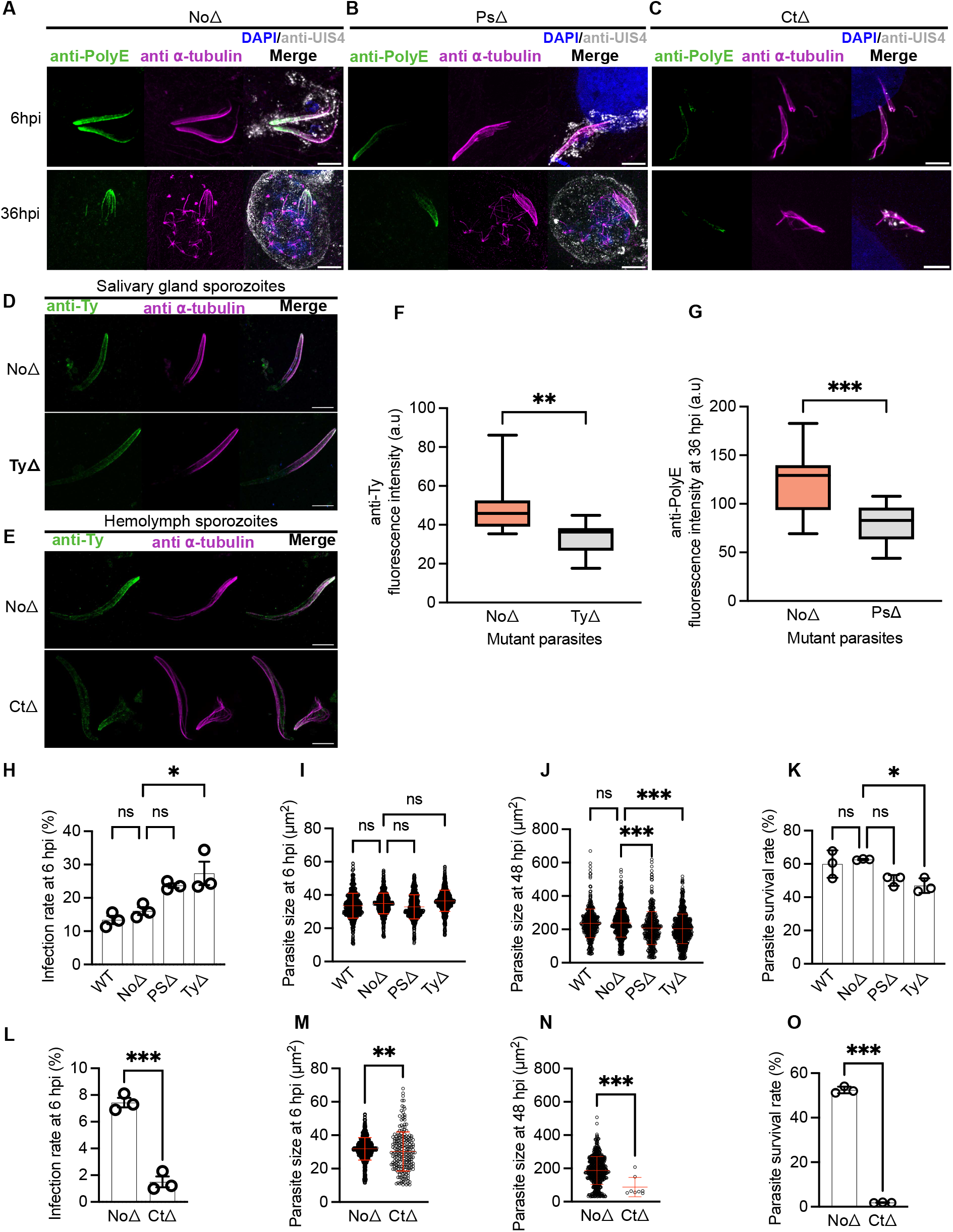
Liver stage development of *Plasmodium* α1-tubulin mutant parasites. **(A–C)** Maximum-intensity projections of confocal z-stacks acquired by PS-ExM from infected HeLa cells at 6 and 36 hours post-infection (hpi). **(A)** NoΔ control parasites derived from salivary gland sporozoites. **(B)** PsΔ parasites derived from salivary gland sporozoites. **(C)** CtΔ parasites derived from hemolymph sporozoites. Samples were labelled with anti-PolyE (green) and anti-α-tubulin (magenta) antibodies. Nuclei were stained with DAPI (blue), and the PVM was labelled with an anti-UIS4 antibody (grey). Scale bars, 10 µm. **(D,E)** Maximum-intensity projections of PS-ExM confocal z-stacks of sporozoites. **(D)** NoΔ and TyΔ salivary gland sporozoites. **(E)** NoΔ and CtΔ hemolymph sporozoites. Samples were labelled with anti-Ty (green) and anti-α-tubulin (magenta) antibodies. Scale bars, 10 µm. **(F, G)** Quantification of antibody-signal intensities. Box-and-whisker plots show the median, interquartile range and minimum-to-maximum values. **(F)** Anti-Ty antibody-signal intensities in NoΔ and TyΔ salivary gland sporozoites. The mean intensity was significantly lower in TyΔ sporozoites (33.4 a.u.) than in NoΔ controls (49.6 a.u.). **(G)** Anti-PolyE antibody-signal intensities in NoΔ and PsΔ liver stage parasites at 36 hpi. The mean intensity was significantly lower in PsΔ parasites (79.9 a.u.) than in NoΔ controls (121 a.u.). Fifteen parasites per line were analyzed across three independent experiments. **(H–K)** Liver stage development following infection with salivary gland sporozoites of the WT, NoΔ, PsΔ and TyΔ lines. **(H)** Infection rates at 6 hpi, calculated as the number of intracellular parasites detected by quantitative microscopy relative to the 20,000 sporozoites used for infection. PsΔ showed no reduction relative to NoΔ, whereas TyΔ showed a modest increase. **(I, J)** Parasite areas at 6 **(I)** and 48 hpi **(J)**. At 48 hpi, PsΔ and TyΔ parasites had mean areas of 208.16 and 203.7 µm², respectively, and were significantly smaller than NoΔ parasites (237.9 µm²). WT and NoΔ parasites were comparable in size. **(K)** Parasite survival between 6 and 48 hpi, normalized to the corresponding parasite number at 6 hpi. Survival at 48 hpi was 60% for WT, 63% for NoΔ, 50% for PsΔ and 47% for TyΔ. **(L–O)** Liver stage development following infection with NoΔ or CtΔ hemolymph sporozoites. **(L)** Infection rates at 6 hpi. CtΔ sporozoites showed a markedly reduced ability to establish intracellular infection. **(M, N)** Parasite areas at 6 **(M)** and 48 hpi **(N)**. The small number of CtΔ parasites persisting at 48 hpi had a mean area of 88.1 µm² and remained significantly smaller than NoΔ controls (187.4 µm²). **(O)** Parasite survival between 6 and 48 hpi, normalized to the corresponding parasite number at 6 hpi. CtΔ survival was reduced to 2%, compared with 52% for NoΔ hemolymph controls. These results indicate severely impaired establishment, survival and growth of CtΔ parasites during liver stage development. PsΔ contains an E445A substitution at a predicted polyglutamylation-associated residue; TyΔ lacks the C-terminal YEADY motif associated with the anti-Ty antibody epitope; CtΔ lacks the α1-tubulin C-terminal tail; and NoΔ is a construct control without a coding-sequence mutation. WT, wild-type parasite. In **(I, J, M, N)**, each dot represents one parasite. In **(H, K, L, O)**, each circle represents the mean of one independent experiment. Summary data are presented as mean ± SD. Statistical analyses for **(H–K)** were performed using one-way ANOVA followed by Dunnett’s multiple-comparisons test against NoΔ. Statistical analyses for **(F, G, L–O)** were performed using two-tailed unpaired *t*-tests. All experiments were performed in three independent biological replicates. *p ≤ 0.05; **p ≤ 0.01; ***p ≤ 0.001.

To examine whether the C-terminal mutations affected antibody recognition, we performed PS-ExM on sporozoites and labelled them with anti-Ty antibodies. Anti-Ty antibody signals remained detectable in both TyΔ salivary gland sporozoites and CtΔ hemolymph sporozoites (Figure 5D and E). Quantification revealed significantly reduced anti-Ty fluorescence in TyΔ salivary gland sporozoites compared with stage-matched NoΔ controls (Figure 5F).

The persistence of anti-PolyE and anti-Ty antibody signals in the PsΔ and TyΔ mutants indicates that antibody recognition was not completely abolished by the respective mutations. These residual signals could reflect recognition of adjacent epitopes, modification of alternative residues, detection of another tubulin isoform, including β-tubulin, or antibody cross-reactivity. Their molecular basis therefore remains unresolved. In particular, the detection of anti-Ty signals in the absence of an identifiable canonical tubulin tyrosine ligase (TTL) in the *Plasmodium* genome (28) warrants further investigation.

To assess parasite survival and growth during liver stage development, we performed quantitative microscopy (Figure 5H to O). Both PsΔ and TyΔ mutant parasites infected HeLa cells and progressed through liver stage development (Figure 5B, D, H, I, J and K). This was further confirmed by staining the PVM using anti-UIS4 antibodies (Figure 5A, B and C). Parasite size was measured at 6 hpi and 48 hpi to assess developmental progression. No significant differences were observed at 6 hpi (Figure 5I). However, by 48 hpi, mutant parasites were significantly smaller than controls, indicating impaired growth (Figure 5J). Survival rates of PsΔ and TyΔ mutants were comparable to controls (Figure 5K), with survival rates of 23.7% for PsΔ and 27.3% for TyΔ, exceeding the control. As expected, the CtΔ hemolymph sporozoites were severely impaired in their ability to establish productive liver stage infections with a typical PVM at all examined time points (Figure 5C, M, N and O).

To place our findings in the context of previous work, we compared our mutants with the previously generated *P. berghei* α1-tubulin mutants described by Spreng et al. (2019) (39). These included parasites lacking the α1-tubulin introns (α1 Δintrons), parasites lacking the C-terminal ADY residues (α1 ΔC-term), and α1-tubulin replacement lines with increasing sequence similarity to α2-tubulin, including the α2+++ line. These mutants showed similar phenotypes, including a complete absence of salivary gland sporozoites in α2-tubulin exon swap parasites, mirroring the CtΔ mutant phenotype (Figure 4D and H; Figure 4–figure supplement 1D and E). Together, these results demonstrate that the α1-tubulin C-terminal region contributes to sporozoite development, motility, and salivary gland colonization.

The severe reduction in salivary gland sporozoites in the CtΔ mutant highlights the importance of this region for successful parasite development and transmission through the mosquito. Future studies should determine the molecular identities underlying the observed anti-PolyE and anti-Ty signals and identify the parasite or host enzymes responsible for their developmental regulation. Biochemical validation combined with targeted genetic manipulation of candidate tubulin-modifying enzymes will be required to establish the functions of individual tubulin modifications in *Plasmodium*.

## Discussion

Our study provides the first in-depth characterization of microtubule architecture and dynamics during the *Plasmodium* liver stage. Using tubulin trackers and tubulin antibodies in HeLa cells and primary mouse hepatocytes, we identified two major microtubule populations: a prominent subpellicular network we termed the LSPMB, and the polymerizing hemi-spindles involved in nuclear division. The LSPMB appears to originate from the SSPMTs and persists until late schizogony. These observations align with earlier electron microscopy studies by Meis et al. (70), who reported the presence of subpellicular microtubules and inner membranes in liver stage parasites up to the point of mitotic spindle formation.

Microtubule behavior and function can be regulated by tubulin PTMs. Sequence analysis showed that *P. berghei* α1-tubulin contains C-terminal residues associated with polyglutamylation and tyrosination in other eukaryotes (Figure 4). Anti-PolyE and anti-Ty antibodies detected signals along sporozoite SSPMTs, consistent with the presence of epitopes associated with these modifications (Figure 3), although their precise molecular identities remain to be established biochemically. The coexistence of different tubulin PTMs on the same microtubule populations has been reported previously (24). In canonical eukaryotic systems, tyrosinated microtubules are generally associated with dynamic populations, whereas detyrosinated microtubules tend to be more stable.

No canonical TTL homolog has been identified in the *Plasmodium* genome (29), leaving the molecular basis of the developmentally regulated anti-Ty staining pattern unresolved. Anti-Ty signals were prominent in invasive sporozoites, became undetectable during much of intracellular liver stage development, and reappeared during the cytomere stage and merozoite formation (Figure 3B). These observations reveal developmentally regulated changes in antibody recognition but do not establish enzymatic tyrosination– detyrosination cycling. The observed pattern could reflect an uncharacterized parasite enzyme, the involvement of host factors, or changes in epitope accessibility or tubulin organization. Nevertheless, its temporal regulation suggests that alterations in the α-tubulin C-terminal region accompany the adaptation of the microtubule cytoskeleton to different stages of the parasite life cycle.

Polyglutamylation introduces negative charges on the tubulin surface that can enhance interactions with microtubule-associated proteins (MAPs) and is commonly associated with stable microtubule populations (71–73). This modification has also been implicated in merozoite biology and intracellular trafficking during *Plasmodium* blood stage development (28). In sporozoites, the concurrent detection of anti-Ty and anti-PolyE signals may indicate a microtubule population combining structural stability with the capacity for regulated interactions during migration through the mosquito and vertebrate host. Following hepatocyte invasion, the anti-Ty signal became undetectable, whereas the anti-PolyE signal persisted.

Importantly, however, the progressive shrinkage and oryzalin sensitivity of the anti-PolyE-positive LSPMB indicate that this structure undergoes extensive remodeling during liver stage development. Rather, the altered pattern of PTM-associated antibody signals may accompany a functional remodeling of the SSPMTs as the slender, motile sporozoite transforms into a rounded and rapidly replicating liver stage parasite. The liver stage is one of the fastest-replicating phases in the *Plasmodium* life cycle, with replication rates among the highest observed in eukaryotic cells (74,75). During this phase, the parasite synthesizes proteins de novo and hijacks host cell lipids and proteins for membrane biogenesis (76–79). We propose that the parasite brings its SSPMTs into hepatocytes to be repurposed as the LSPMB (Figure 6). Their progressive disassembly could provide a readily available pool of tubulin for the extensive microtubule polymerization required during schizogony, while potentially reducing the energetic cost of de novo tubulin synthesis.

**Figure 6.**
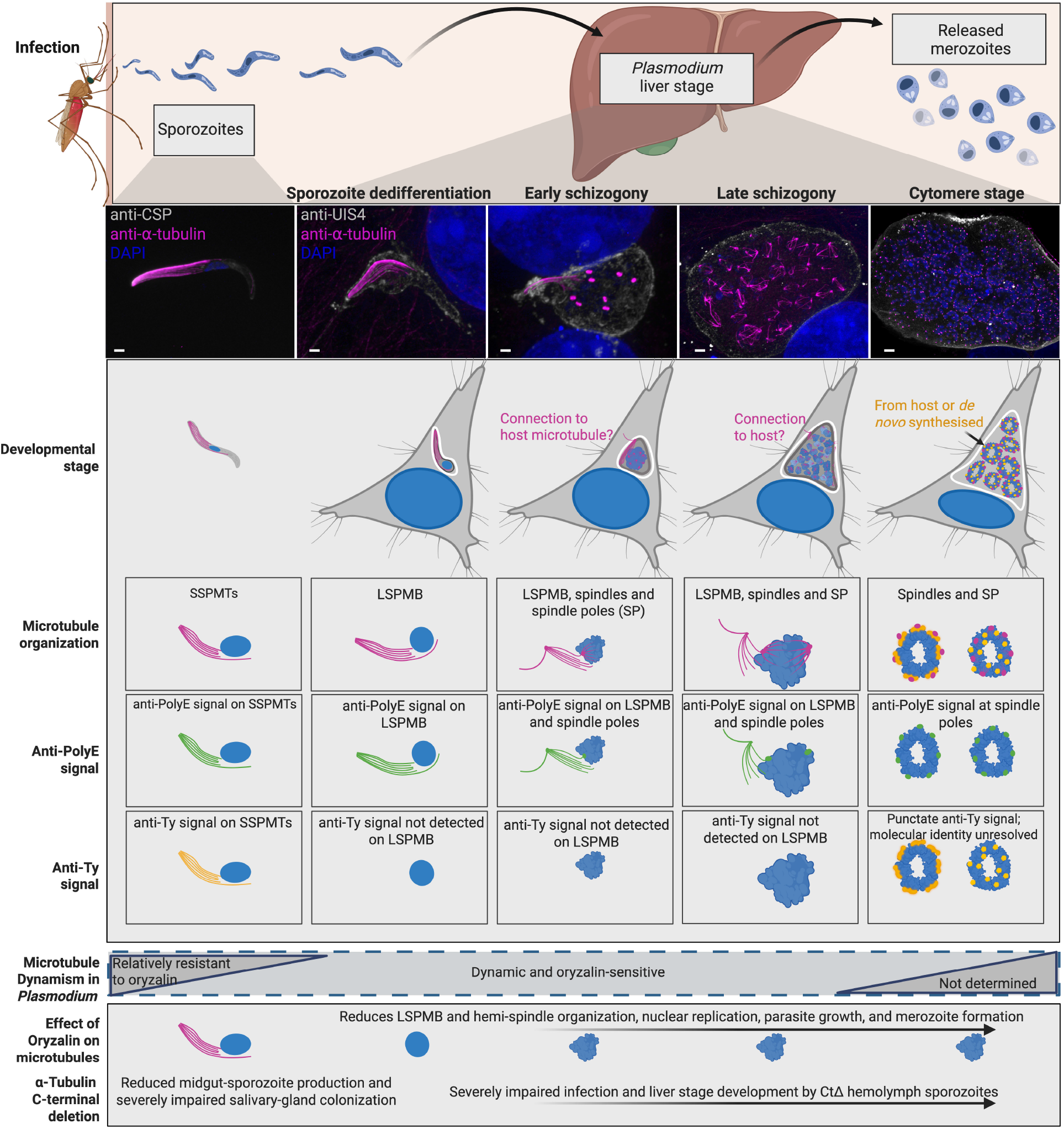
Summary model of microtubule organization during *Plasmodium berghei* sporozoite and liver stage development. The model summarizes the principal observations of this study from salivary gland sporozoites through liver stage development and merozoite formation. Representative maximum-intensity projections show α-tubulin labelled with an anti-α-tubulin antibody (magenta), nuclei stained with DAPI (blue), and either the sporozoite surface labelled with an anti-CSP antibody or the parasitophorous vacuole membrane labelled with an anti-UIS4 antibody (grey). Scale bars, 10 µm. Following hepatocyte invasion, the sporozoite subpellicular microtubules (SSPMTs) persist and remodel into the liver stage parasite microtubule bundle (LSPMB). During schizogony, the LSPMB progressively decreases in size, and individual LSPMB filaments are transiently found in close association with hemi-spindle poles. At the cytomere stage, the LSPMB is no longer detected, while microtubules associated with spindle poles and developing merozoites are abundant. Anti-PolyE antibody signals are detected on the SSPMTs, the LSPMB and structures associated with hemi-spindle poles. Anti-Ty antibody signals are detected on sporozoite SSPMTs but are not detected on the LSPMB during early and late schizogony. Punctate anti-Ty signals reappear during the cytomere stage, although the molecular identity of the labelled structures remains unresolved. These antibody-staining patterns should not be interpreted as definitive biochemical identification of the corresponding tubulin modifications. Sporozoite SSPMTs are relatively resistant to oryzalin, whereas liver stage microtubule structures are oryzalin-sensitive. Oryzalin treatment reduces LSPMB and hemi-spindle organization and impairs nuclear replication, parasite growth and merozoite formation. Mutational analysis further indicates that the α1-tubulin C-terminal region contributes to mosquito stage development and liver stage infection. Deletion of the α1-tubulin C-terminal tail reduces midgut sporozoite production, severely impairs salivary gland colonization, and markedly reduces the ability of hemolymph sporozoites to establish and maintain liver stage infection.

Consistent with such a role, LSPMB filaments were observed spatially associated with hemi-spindle poles. These spatial associations were relatively rare in fixed samples but occurred more frequently during live imaging, suggesting that they are transient. Together, these observations raise the possibility that tubulin originating from the LSPMB is redistributed to support hemi-spindle formation during nuclear replication. EB1–eGFP was detected in association with both the LSPMB and hemi-spindles during liver stage development. EB1 is a conserved microtubule plus-end-binding protein that regulates microtubule growth and recruits additional microtubule-associated proteins in other systems. Although EB1 expression in *Plasmodium* was previously reported primarily during sexual blood stages (53–55), we also detected EB1–eGFP during mosquito and liver stages. Its localization is consistent with a potential involvement in parasite microtubule organization, although functional studies will be required to determine whether EB1 contributes to LSPMB remodeling or spindle formation.

To investigate the functional contribution of the α1-tubulin C-terminal region, we generated *P. berghei* mutants carrying targeted alterations in residues associated with PTM-related antibody epitopes. Unexpectedly, anti-PolyE and anti-Ty antibody signals remained detectable after mutation of the predicted polyglutamylation-associated residue or deletion of the C-terminal YEADY motif, respectively, although signal intensities were reduced relative to the corresponding controls.

The residual antibody signals could reflect recognition of adjacent epitopes, modification of alternative residues, detection of another tubulin isoform, or antibody cross-reactivity. Their molecular basis therefore remains unresolved. Mass spectrometry-based mapping will be required to establish which tubulin residues are modified in the different parasite lines and whether alternative modifications occur at neighboring sites. Importantly, the CtΔ mutant produced very few typical salivary gland sporozoites and consequently showed severely impaired liver stage establishment, survival and growth, demonstrating an important contribution of the α1-tubulin C-terminal region to sporozoite development and transmission.

Previous studies have also demonstrated partial redundancy between α1- and α2-tubulin isoforms (38,39,80). However, the complete C-terminal deletion mutant produced very few typical salivary gland sporozoites and showed severely impaired liver stage establishment, survival and growth. These findings are consistent with Spreng et al. (2019) (39), who reported defective sporozoite formation, altered microtubule numbers, and impaired motility in α1-tubulin mutants lacking C-terminal sequences.

In conclusion, our study reveals a previously unrecognized complexity in the microtubule biology of the *Plasmodium* liver stage. We propose that the stable subpellicular microtubules carried into the hepatocyte by the sporozoite are repurposed into a dynamic LSPMB that is progressively remodeled during liver stage development and may contribute tubulin to the rapidly expanding microtubule network required for schizogony. This remodeling is accompanied by developmentally regulated changes in PTM-associated antibody signals and involves microtubule-associated proteins such as EB1. The pronounced phenotypes caused by modification of the α1-tubulin C-terminal region further demonstrate the importance of tubulin regulation for parasite development and transmission. Determining the molecular identities underlying these antibody signals and identifying the enzymes potentially responsible for the corresponding modifications should provide important insights into *Plasmodium* biology and may reveal new targets for therapeutic intervention.

## Methods

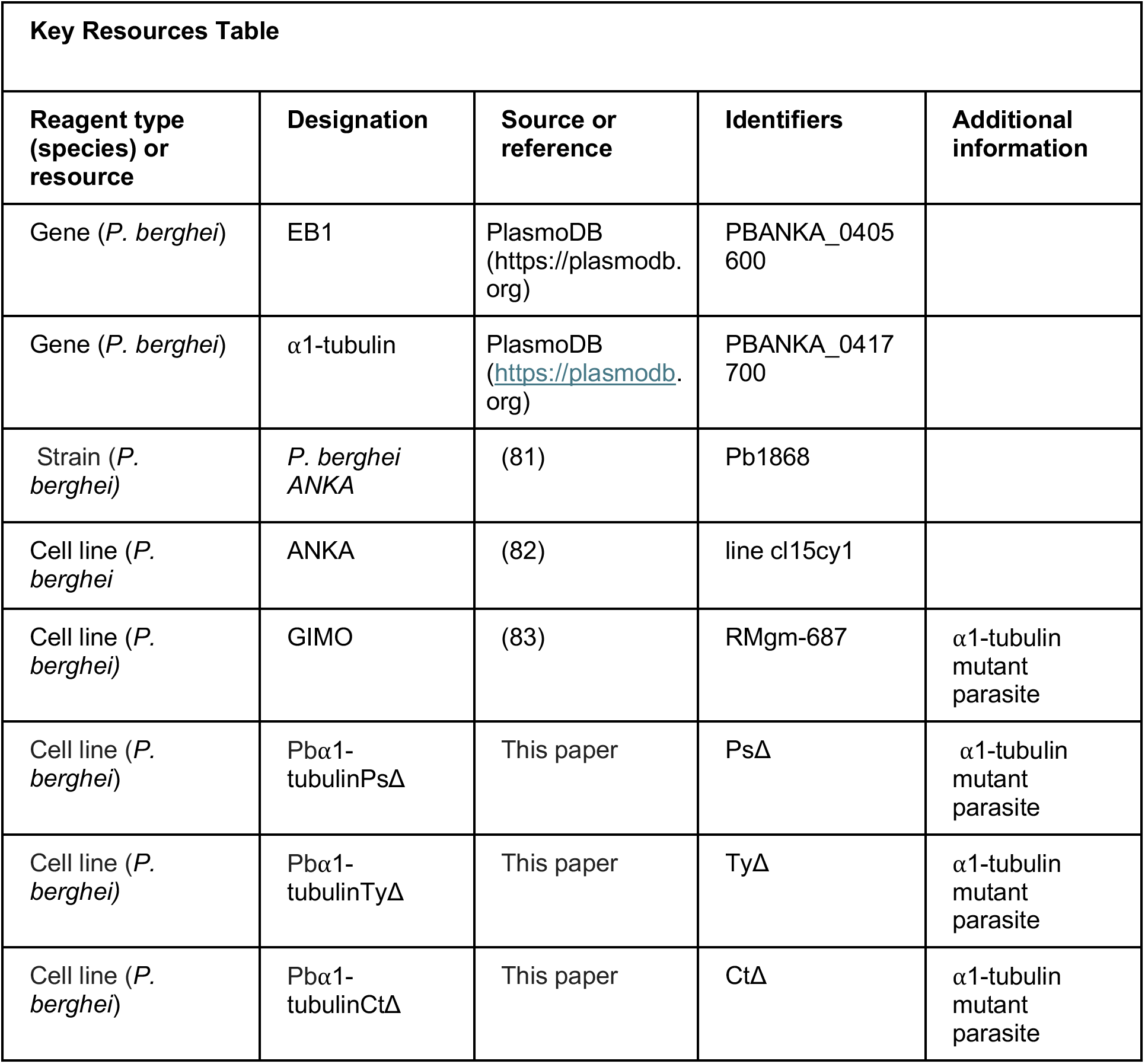
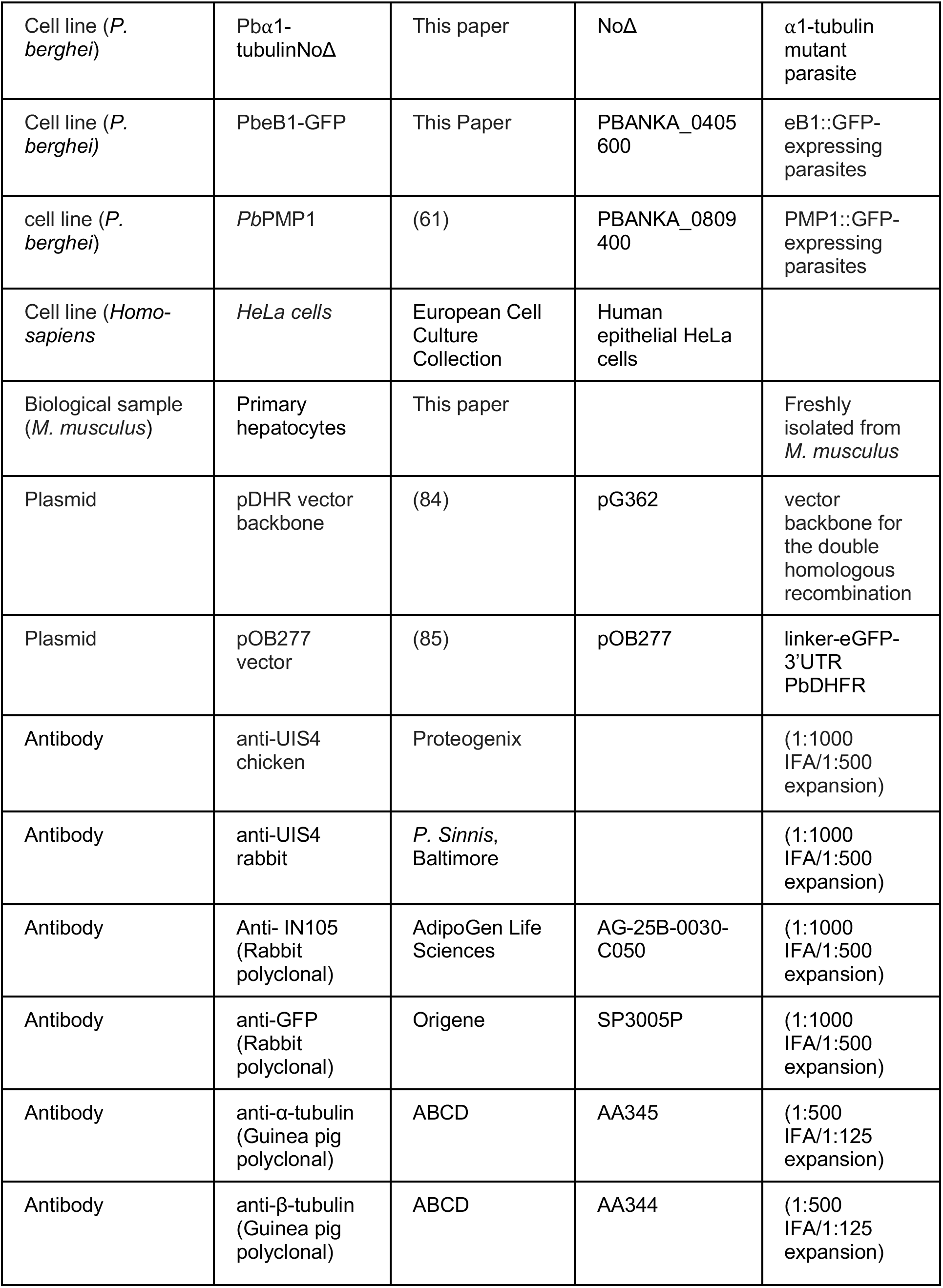
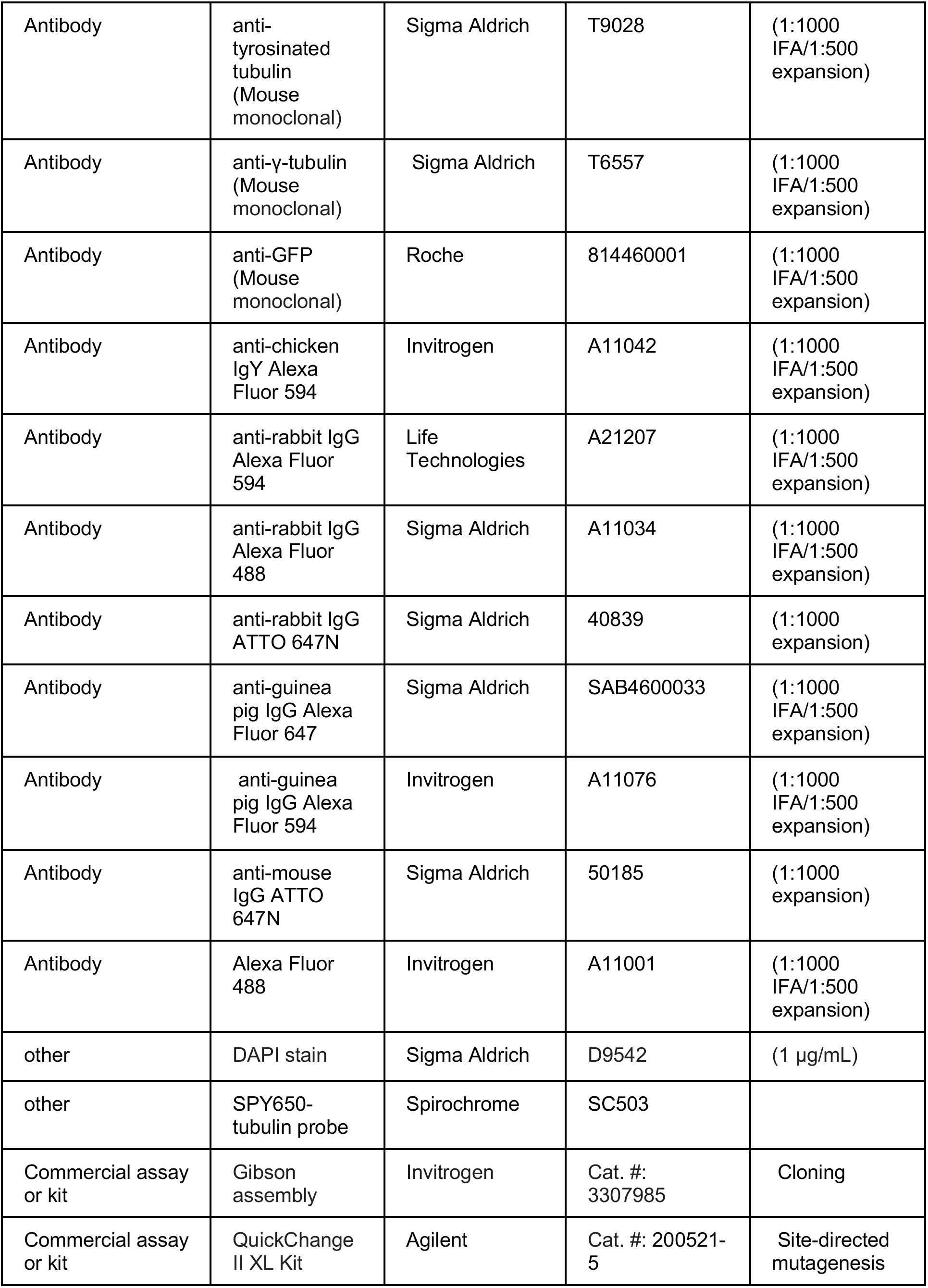
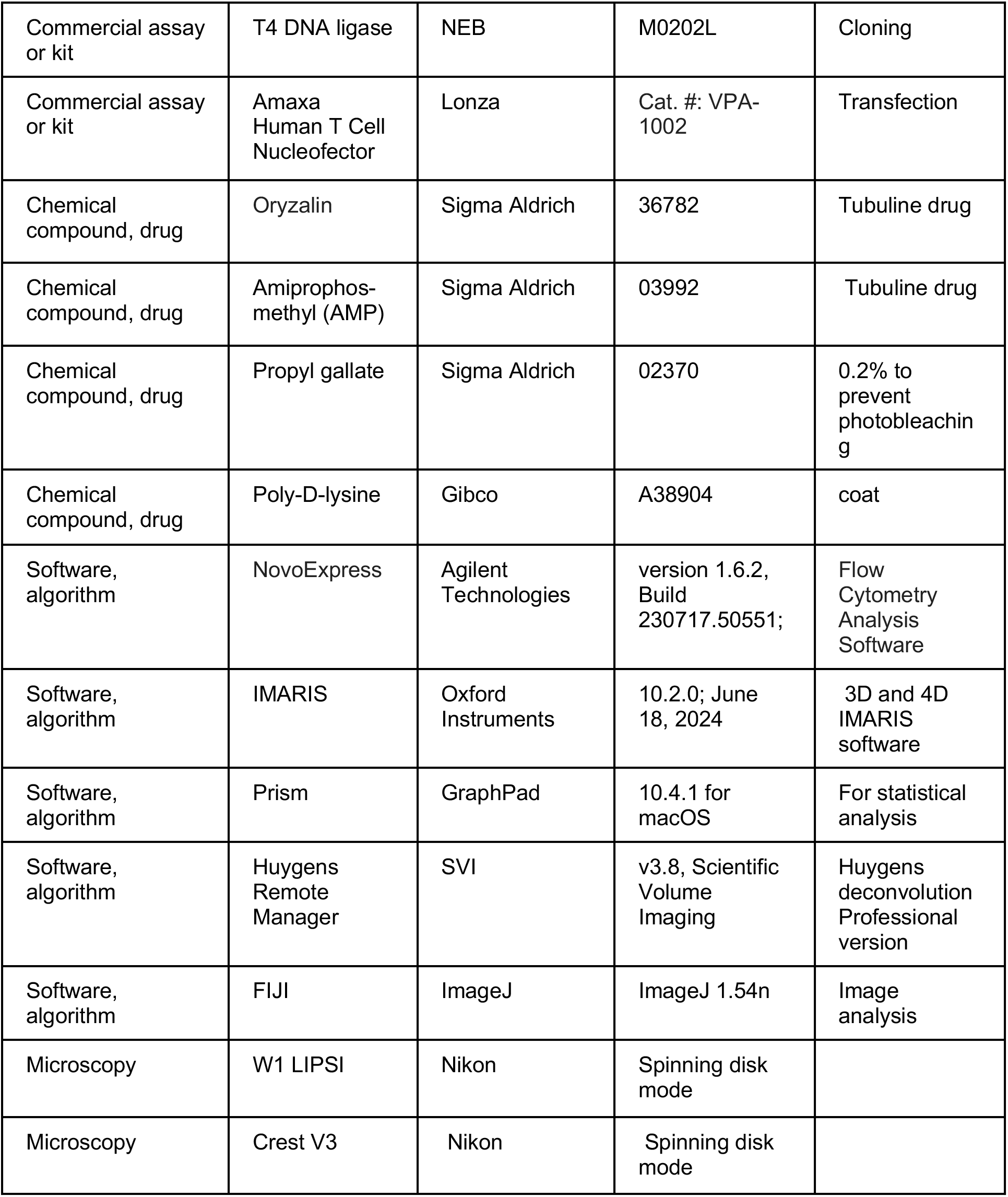

### Ethics statement

All experimental procedures were performed in full compliance with the Swiss Animal Protection Act (TSchG) and were approved by the Canton of Bern’s Animal Experimentation Commission (Authorization BE118/22). Personnel involved in animal studies held Federation of European Laboratory Animal Science Associations (FELASA)-accredited certifications. Female C57BL/6 and BALB/c mice of 6 to 10 weeks old and 20-30 g at infection were sourced from Janvier Labs (France) or bred in the animal facility at the University of Bern’s Institute of Cell Biology. Animals were maintained under standard housing conditions with daily health monitoring by trained staff.

### *P. berghei* lines used in this study and the generation of transgenic parasites

The *P. berghei* ANKA parasite line was utilized for both murine and cellular infection models. To generate the α1-tubulin C-terminal mutant and EB1::eGFP parasite lines, the C-terminal coding and 3’ untranslated regions (UTRs) were cloned into the double homologous recombination vector pDHR, which contains the 3’UTR PbDHFR, the ef1*α* promoter region, the selectable markers hdhfr::yfcu, and additional 3’UTR PbDHFR sequences. The pDHR vector backbone originated from the plasmid pG362 (kindly provided by the Waters lab) (84).

First, the pG362 plasmid was digested with SacII and PstI to remove the AID, selectable marker, and GFP sequences. The amplified product (linker-eGFP-3’UTR PbDHFR) from the pOB277 vector was then cloned using T4 DNA ligase (NEB, M0202L) (85). For subsequent cloning, restriction sites were added to the forward (KpnI and ApaI) and reverse (EcoRI and SmaI) primers. The plasmid was then digested with EcoRI and SmaI to clone the amplified product: the ef1α promoter region-hdhfr::yfcu marker (a fusion gene of the positive selection marker human dihydrofolate reductase (hdhfr) and the negative selection marker yeast cytosine deaminase-uridyl phosphoribosyl transferase (yfcu)) 3′UTR PbDHFR from the genomic DNA of the *P. berghei* GIMO parasites (RMgm-687) parental line (83).

The complete *α*1-tubulin gene sequence (PBANKA_0417700), including 5’ and 3’ UTRs, was retrieved from PlasmoDB (https://plasmodb.org). The linker-eGFP region of the pDHR vector was removed using ApaI and NotI restriction enzymes. The 5’UTR of the *α*1-tubulin gene was then subcloned into the digested pDHR using Gibson assembly. The resulting vector was digested with SmaI, and the PbHsp70 promoter and mCherry regions from RMgm-687, along with the 3’UTR of the *α*1-tubulin gene, were subcloned into the vector at the ApaI and SmaI restriction sites, respectively, using Gibson assembly.

Site-directed mutagenesis (QuickChange II XL Kit) was used to introduce the C-terminal modifications, with primer sequences listed in **Appendix 1 – Tables 2 and 3**. The final plasmid was linearized with ScaI before being transfected into the PbWT line (*P. berghei* ANKA line cl15cy1) (82). Similarly, the end-binding protein EB1 (PBANKA_0405600) with an eGFP tag was engineered using identical cloning strategies. The 5’ and 3’ UTRs were retrieved from PlasmoDB (https://plasmodb.org) and subcloned into the pDHR vector at the ApaI and SmaI restriction sites, respectively, using Gibson assembly. The final vector was linearized with NdeI before being transfected into the Pb1868 line (constitutively expressing Hsp70-mCherry-ef1α-NLuc) (81). Primer sequences are listed in **Appendix 1 – Table 1**.

Vector transfections were performed in schizont cultures as previously described (64,86). Briefly, *P. berghei* schizonts were cultured and purified (64,86,87). Electroporation of schizonts from the respective parasite lines was performed using the Amaxa® Human T Cell Nucleofector® system with the U33 program, followed by intravenous injection into naïve mice. The construct was integrated by double-crossover homologous recombination at the locus of interest. Pyrimethamine drug selection was administered via drinking water, and parasitemia was monitored daily from day 7 post-infection until mice tested positive (86).

### Parasitemia measurement and cloning of transgenic parasite lines

To measure the parasitemia ratio of infected mice, approximately 2 μL of blood from the tail vein of each infected mouse was added to 500 μL of PBS and analyzed by flow cytometry using the NovoCyte (Agilent, 2023) as described (88). The data were analyzed using NovoExpress Flow Cytometry Analysis Software (version 1.6.2, Build 230717.50551; Agilent Technologies).

To clone the different engineered transgenic parasite lines, mice were infected with the transgenic parasites. Three days post-infection, parasitemia was assessed, and when parasitemia reached at least 1%, blood was collected from the infected mouse via heart puncture following euthanasia as described (64). The collected blood was passaged into a new naïve mouse. When parasitemia in the newly infected mouse reached 1-2%, 2 μL of tail vein blood was collected and mixed with RPMI-1640. Maintaining low parasitemia was crucial to prevent both double erythrocyte infection by *P. berghei* and gametocyte formation. Single *P. berghei*-infected erythrocytes were then sorted into individual wells of 96-well plates (Cellvis) containing 200 μL of RPMI-1640 supplemented with 10% FCS using a BD FACSDiscover S8 with CellView. Supplementing the medium with 10% FCS allowed the sorted infected erythrocytes to recover from the sorting process. Five mice were then intravenously infected with a single sorted infected erythrocyte each. The mice underwent drug selection via drinking water, and parasitemia was monitored daily from day 7 post-infection until parasites were detected. Clonality of the parasites in positive mice was verified by integration PCR using primers listed in **Appendix 1 – Table 1, 2, and 3**.

### Cell culture

Human epithelial HeLa cells (European Cell Culture Collection) were maintained in Minimum Essential Medium (MEM; 1-31F01-I BioConcept) supplemented with Earle’s salts and 10% heat-inactivated fetal bovine serum (FBS; Sigma-Aldrich), hereafter referred to as complete MEM (cMEM). Primary hepatocytes were cultured in William’s Medium E (WME) complete medium. All cells were incubated at 37°C under 5% CO*₂*. For infection assays, 40,000 HeLa cells were seeded onto coverslips in 24-well plates and allowed to adhere overnight. Cells were then infected with *P. berghei* sporozoites and maintained in cMEM containing 2.5 µg/mL amphotericin B (PAA Laboratories, P11-001) to prevent fungal contamination. Medium was exchanged every 24 hours.

In case of microtubule drug treatment, the cells were incubated with the drug continuously for the duration of the experiments. To maintain consistent drug pressure, the culture medium was replaced every 24 hours with fresh medium containing the specified concentration of the drug. For the reversibility experiments, the drug-containing medium was removed after the initial treatment period, and the cells were washed five times with drug-free medium. They were then incubated in fresh medium without the drug for the remainder of the experiment to assess recovery.

### Primary hepatocyte isolation

The mouse primary hepatocytes were isolated according to the method described (89). Briefly, the cells were isolated from 12 to 16 week old Balb/c or C57BL/6 mice using a two-step perfusion technique (76,89,90). Mice were euthanized via CO*₂* inhalation (1–2 minutes), and after securing the animal in a supine position, a midline abdominal incision was made to expose the portal vein or inferior vena cava (IVC). A peristaltic pump was pre-rinsed with 70% ethanol and HEPES buffer, then set to 50 for initial cannulation of the portal vein (or IVC) using a Teflon-coated needle. Upon successful insertion, the flow was increased to maximum (999), and the opposing vein was severed to allow perfusion with HEPES buffer for 5 minutes, during which periodic clamping (10 seconds per minute) of the cut vein was performed. The tubing was subsequently changed to a collagenase solution, and perfusion was held for 7 minutes with repeated clamping. The liver was excised, transferred to a collagenase-containing well, and mechanically dissociated using filter tips and gentle shaking with forceps. Cell suspensions were sequentially collected, pooled, and washed with William’s Medium E (WME) by centrifugation at 50 g for 2 minutes. The pellet was resuspended in WME, optionally passed through a 70 µm nylon strainer, and washed twice more. The final pellet was resuspended in WME complete medium (WME supplemented with L-glutamine, penicillin-streptomycin, amphotericin B, insulin-transferrin, and aprotinin), and cells were counted and seeded at densities appropriate for T25 flasks, GB dishes, or 24-well plates. After initial adhesion (1 hour), culture medium was supplemented or exchanged after 2–4 hours.

### Immunofluorescence assay (IFA) of fixed primary hepatocytes and HeLa cells

Immunofluorescence was carried out as previously described (45,79,91). Cells were seeded onto coverslips in culture 24-well plates and allowed to adhere overnight before infection with 20,000 sporozoites. Infection was maintained for 6, 24, 36, 48 and 56 hours post-infection (hpi), after which cells were fixed using cold methanol at −20°C for 15 minutes and subsequently washed twice with phosphate-buffered saline (PBS). Blocking was performed by incubating cells in 10% FCS in PBS for 20 minutes at room temperature to reduce non-specific binding. Primary antibody staining was carried out by incubating cells in 10% FCS diluted in PBS for 1 hour at room temperature, followed by three washes with PBS. Secondary antibody incubation was performed in 10% FCS as well for 1 hour at room temperature in the dark. The antibodies used are listed in the enumerated section below. After three additional washes with PBS, nuclei were counterstained with DAPI (4′,6-diamidino-2-phenylindole). Coverslips were then mounted onto slides with DAKO mounting medium and fluorescence imaging was performed using either Nikon W1 or LIPSI and Crest V3 confocal microscope equipped with filters for DAPI, AF488, AF594, and Cy5. Single-stain controls were carried out for every staining to ensure that bleed-through was negligible.

### Sporozoite isolation from mosquito salivary glands and hemolymph

*Anopheles mosquitoes* were fed on *Plasmodium*-infected mice exhibiting a gametocytaemia of 0.2 to 1% and the sporozoite isolations were performed from day 18 post-infection as previously described (50,64). For salivary gland sporozoites, mosquitoes were anesthetized with chloroform, sterilized in 70% ethanol, and rinsed twice in PBS. The mosquitoes were then dissected, and salivary glands were extracted under a binocular microscope in Iscove’s Modified Dulbecco’s Medium (IMDM) pooled in 50 µL of ice-cold IMDM and mechanically homogenized using a pestle (Sigma-Aldrich, Z359947) powered with cordless motor (Sigma-Aldrich, Z359971) as described (50,64). For hemolymph sporozoite extraction, glass capillaries were heat-pulled to fine tips. The abdominal terminal segment of the mosquito was excised, and hemolymph was flushed from the thorax by PBS perfusion via capillary insertion (68,92). The extracted hemolymphs were centrifuged (11,000 × g, 1 min), and pellets were resuspended in PBS or IMDM. Sporozoite quantification was performed using a Neubauer hemacytometer. Midgut, hemolymph, or salivary gland suspensions were mixed 1:1 with RPMI-1640 supplemented with 10% FCS, and sporozoites within four corner grids were counted. Total yields (sporozoites/mL and per mosquito) were calculated based on dilution factors and dissection numbers.

### Sporozoites IFA and gliding motility assay

To perform the sporozoite immunofluorescence assay (IFA), 12 mm cover glasses (0111530, Paul Marienfeld, Germany) were coated with 0.1 mg/mL poly-D-lysine (A38904-01, Gibco, USA) (45,93) and incubated overnight at room temperature. The following day, sporozoites were isolated from mosquito midguts, hemolymph, or salivary glands as described above. The extracted sporozoites were suspended in RPMI-1640 medium to a final volume of 200 µL in an Eppendorf tube and centrifuged at 1000 rpm for 3 min. The pellet was fixed by resuspension in 200 µL of cold methanol and incubated for 15 min at −20°C. After fixation, sporozoites were washed twice with PBS (1000 rpm centrifugation) and transferred onto poly-D-lysine-coated cover glasses in a 24-well plate (400024, Bioswisstec, Switzerland). The plate was incubated for 30 min at room temperature to facilitate sporozoite adhesion before proceeding with either the IFA or expansion microscopy protocol.

For the gliding motility assay, 5 µL of the sporozoite suspension in RPMI-1640 (supplemented with 10% FCS) was added to 96-well glass-bottom plates (P96-1.5H-N, Cellvis, USA) (45,65,66). The plate was centrifuged (1000 rpm, 1 min) and immediately live imaged using a Leica DM5500 widefield fluorescence microscope.

### Expansion microscopy

Expansion microscopy (including PS-ExM and U-ExM when mentioned) was performed as previously established (45–47). Briefly, infected cells (HeLa cells, primary hepatocytes and sporozoites) were seeded on 13 mm coverslips in 24-well plates and fixed with either 4% paraformaldehyde (PFA; F8775, Sigma) or cold methanol. The cells were anchored by incubation in PBS containing 0.7% formaldehyde and 1% acrylamide at 37°C for 2 hours. Gelation was then performed by adding 5 µL each of 10% tetramethylethylenediamine (TEMED; 17919, Thermo Fisher) and 10% ammonium persulfate (APS) to 90 µL of monomer solution containing 19% sodium acrylate (408220, Sigma), 10% acrylamide (A4058, Sigma), and 0.1% N, N’-methylenebisacrylamide (M1533, Sigma). A 35 µL volume of monomer solution was dispensed onto parafilm in a humidity chamber, and coverslips containing the anchored cells were inverted onto droplets. Following 5 minutes incubation on ice to improve gel penetration, gels were polymerized at 37°C for 30 minutes. Gel-embedded samples were transferred to denaturation buffer (200 mM SDS, 200 mM NaCl, 50 mM Tris-HCl, pH 9) in 6-well plates and agitated (200 rpm) at room temperature for 15 minutes, followed by incubation in fresh buffer at 95°C for 1 hour with gentle shaking. Gel expansion was performed in autoclaved MilliQ water (50 mL) for 1 hour with optional buffer exchange. For immunostaining, gels were incubated with primary antibodies in 2% BSA for 2 hours at 37°C, washed with PBS-Tween 0.1%, then labelled with secondary antibodies plus 1 µg/mL DAPI for nuclear staining in 2% BSA for 2 hours at 37°C. For experiments involving PS-ExM, cell immunofluorescence staining was performed following the established protocol (46) immediately before the gelation process. Final gel expansion was conducted in MilliQ water containing 0.2% propyl gallate (02370, Sigma) to prevent photobleaching. Expanded samples were mounted on poly-D-lysine-coated (A38904, Gibco USA) glass-bottom dishes with cellular surfaces facing downward and imaged with confocal microscopy.

### Antibodies used for IFA and expansion microscopy

The primary antibodies used were: anti-UIS4 chicken (1:1000 IFA/1:500 expansion; Proteogenix), anti-UIS4 rabbit (1:1000 IFA/1:500 expansion; P. Sinnis, Baltimore), polyglutamylation IN105 rabbit (1:1000 IFA/1:500 expansion; AG-25B-0030-C050, AdipoGen Life Sciences) was raised against at least four glutamate residues (GEGE*EE, where * marks the modification site) and is specific for long polyglutamylation side chains. All antibodies were epitope-affinity purified. Anti-GFP rabbit (1:1000 IFA/1:500 expansion; SP3005P, Origene), anti-α-tubulin guinea pig (1:500 IFA/1:125 expansion; AA345, ABCD), anti-β-tubulin guinea pig (1:500 IFA/1:125 expansion; AA344, ABCD), anti-tyrosinated tubulin mouse (anti-Ty, 1:1000 IFA/1:500 expansion; T9028, clone TUB-1A2, Sigma-Aldrich) was a purified monoclonal antibody recognizing C-terminal tyrosine residues on tubulins; it does not react with carboxypeptidase A-treated cells, confirming specificity for the tyrosinated form. anti-α-tubulin tyrosinated rat (1:1000 IFA/1:500 expansion; clone YL1/2, MAB1864-I, EMD Millipore) was a purified immunoglobulin recognizing a linear epitope with a C-terminal aromatic residue flanked by two negatively charged amino acids (as in Gly-Gly-Tyr in tyrosinated tubulin). Anti-γ-tubulin mouse (1:1000 IFA/1:500 expansion; T6557, Sigma Aldrich); anti-GFP mouse (1:1000 IFA/1:500 expansion; 814460001, Roche).

Secondary antibodies included anti-chicken IgY Alexa Fluor 594 (1:1000 IFA/1:500 expansion; A11042, Invitrogen Molecular Probes), anti-rabbit IgG Alexa Fluor 594 (1:1000 IFA/1:500 expansion; A21207, Life Technologies) and 488 (1:1000 IFA/1:500 expansion; A11034, Sigma Aldrich), anti-rabbit IgG ATTO 647N (1:1000 expansion only; 40839, Sigma Aldrich), anti-guinea pig IgG Alexa Fluor 647 (1:1000 IFA/1:500 expansion; SAB4600033, Sigma Aldrich) and 594 (1:1000 IFA/1:500 expansion; A11076, Invitrogen Molecular Probes), and anti-mouse IgG ATTO 647N (1:1000 expansion only; 50185, Sigma Aldrich) and Alexa Fluor 488 (1:1000 IFA/1:500 expansion; A11001, Invitrogen Molecular Probes).

### In vivo assay

To assess the infectivity of parasite mutant sporozoites to hepatocytes, *Anopheles stephensi* mosquitoes infected with different parasite mutants were allowed to feed on anesthetized naïve mice. Three mice per parasite line and five mosquitoes per mouse were used in each replicate. The blood stage infection progression was monitored daily via flow cytometry, starting from the day of infection until mice tested positive.

### Oocyst size measurement

To quantify oocyst size, *Anopheles stephensi* mosquitoes were allowed to feed on anesthetized mice infected with the parasite (Rogerio et al. 2006; Hegge et al., 2009). At 7- and 11-days post-feeding, ten mosquitoes per time point were dissected, and their midguts were extracted and placed in 1× PBS. The midguts were then transferred to a 21-well microscopy slide, covered with a coverslip, and imaged using a Nikon Crest V3 confocal microscope (10× objective). The oocyst area was measured using Fiji software (ImageJ). Each experiment was performed in triplicate for every parasite line.

### Liver stage parasite size measurement

To quantify the parasite size during liver stage development, HeLa cells (40,000 cells/well, triplicate) were seeded in 96-well plates and infected 24 hours later with 20,000 sporozoites per well. After a 2-hour incubation at 37°C, infected cells from each well were washed and redistributed into eight wells across two 96-well plates (4 wells/plate) to optimize cell density for imaging (11,88). Parasite development was analyzed at 6, 24, and 48 hpi using an InCell Analyzer 2000 with 10× objective. The image processing parameters such as kernel size, sensitivity, and size thresholds were optimized in InCell Developer software to distinguish individual parasites from background signals. Parasite growth was assessed based on size measurements, while survival rates were calculated as percentages relative to 6 hpi baseline counts.

### Live-cell imaging of *Plasmodium* microtubule dynamism

HeLa cells were seeded in a 96-well plate and infected the following day with *P. berghei* salivary gland sporozoites. After a 2-hour incubation at 37°C, cells were washed with MEM medium to remove uninfected parasites and residual mosquito salivary gland materials. The infected cells were then reseeded in a dark 24-well glass-bottom plate (Cellvis) and maintained in fresh cMEM at 37°C with 5% CO*₂* (64,94). Prior to live imaging, cells were incubated for 1 hour in cMEM containing the SPY650-tubulin probe (1:1000 dilution; SC503, Spirochrome, Switzerland). Images were acquired using a Nikon W1 LIPSI spinning disk microscope with a 100× objective. Marked infected HeLa cells were imaged at defined time points to generate time-lapse movies capturing tubulin dynamics and parasite development.

### Image processing and data analysis

All the live-cell imaging, expansion microscopy, and IFA images in this study were acquired with Nikon W1 LIPSI and Crest V3, both in spinning disk mode. The microscope images were processed and analyzed using FIJI software, Huygens deconvolution Professional version (Huygens Remote Manager v3.8, Scientific Volume Imaging, The Netherlands), or, when specified, the 3D and 4D IMARIS software (Imaris 10.2.0; June 18, 2024). The 3D/4D image analysis software was used to segment the parasite LSPMB, hemi-spindle, and poles into either isospots or isosurfaces, as previously described (45,64). The counts, volumes, and coefficients of the isospots or isosurfaces were retrieved in .csv files, which were then imported into GraphPad Prism for statistical analysis. All data were analyzed using GraphPad Prism version 10.4.1 for macOS (GraphPad Software, San Diego, California, USA). Statistical comparisons were performed using one-way ANOVA with Dunnett’s multiple comparisons test (*p ≤ 0.05, **p ≤ 0.01, ***p ≤ 0.001). All experiments in the study were carried out in triplicate.

## Acknowledgments

The authors gratefully acknowledge the Microscopy Imaging Center (MIC) at the University of Bern, Switzerland, for their technical support and access to microscopy equipment. K.A. acknowledges the Swiss Confederation for the award of a Government Excellence Scholarship. This work was funded by the Swiss National Science Foundation (SNSF) (grant number 310030_212795) and an Multidisciplinary Center for Infectious Diseases (MCID), University of Bern, Switzerland, grant to Volker Heussler (MA-09).

## Supplementary figures

**Figure 2–figure supplement 1.**
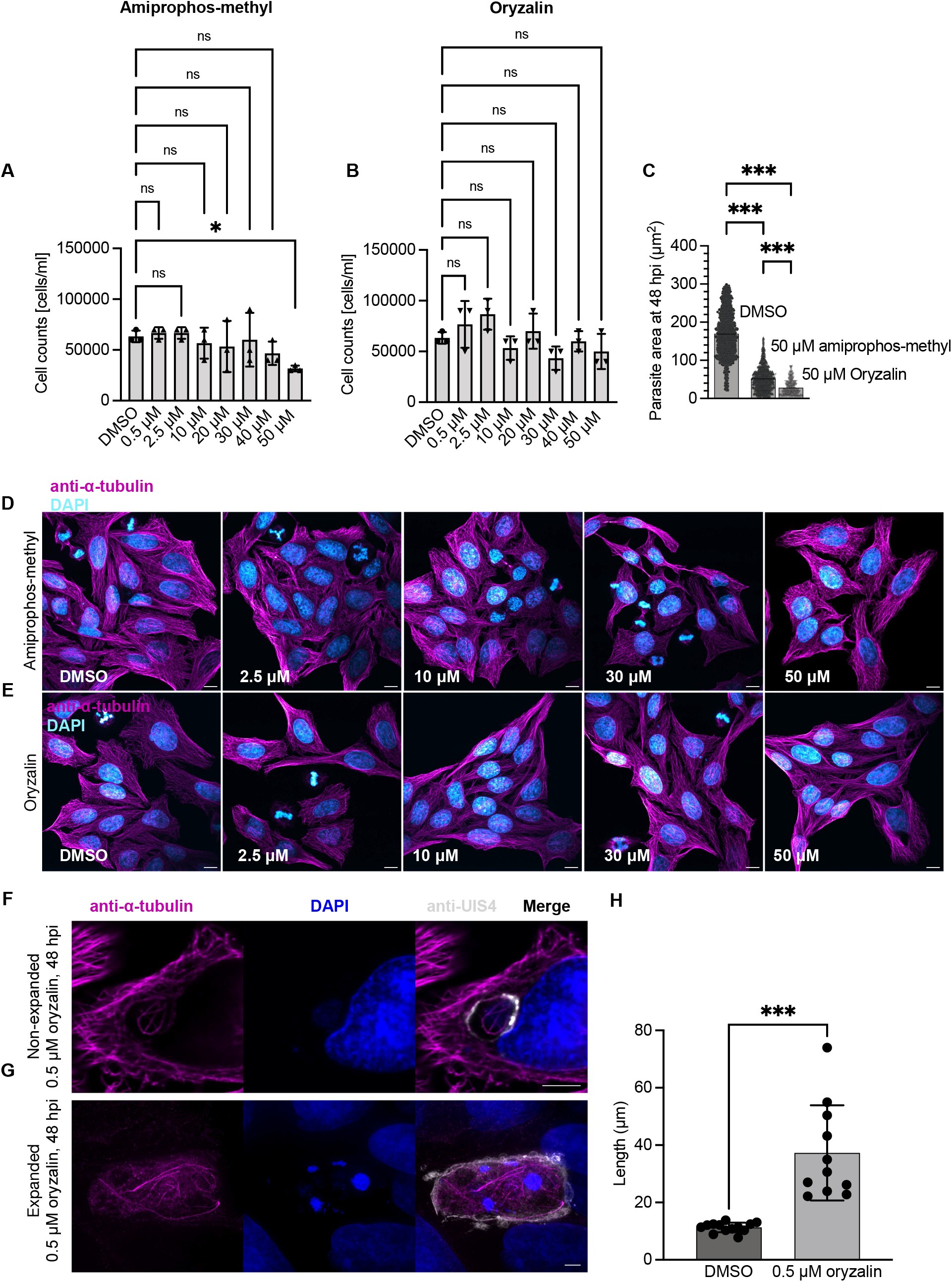
Effects of microtubule-targeting compounds on HeLa cells and *Plasmodium* liver stage development. HeLa cells were seeded in six-well plates and treated the following day with the indicated concentrations of amiprophos-methyl (AMP) or oryzalin. The medium was replaced daily with fresh compound-containing medium. After five days, surviving cells were detached and counted. **(A)** HeLa cell numbers following AMP treatment. A significant reduction relative to the DMSO control was detected only at 50 µM AMP. **(B)** HeLa cell numbers following oryzalin treatment. No significant differences relative to the DMSO control were detected at the tested concentrations. Each point represents one independently treated well from one independent experiment. Data are presented as mean ± SD from three independent experiments. Statistical analysis was performed using one-way ANOVA followed by Dunnett’s multiple-comparisons test against DMSO. **(C)** Parasite area at 48 hpi following treatment with DMSO, 50 µM AMP or 50 µM oryzalin. Oryzalin treatment reduced parasite growth relative to both DMSO and AMP treatment. Each dot represents one parasite. Statistical analysis was performed using one-way ANOVA followed by Tukey’s multiple-comparisons test. **(D, E)** Representative maximum-intensity projections of confocal z-stacks of HeLa cells treated with the indicated concentrations of AMP **(D)** or oryzalin **(E)**. Cells were fixed with cold methanol, microtubules were labelled with an anti-α-tubulin antibody (magenta), and nuclei were stained with DAPI (blue). Scale bars, 5 µm. **(F)** Representative maximum-intensity projections of confocal z-stacks of non-expanded infected HeLa cells treated with 0.5 µM oryzalin and fixed at 48 hpi. Parasite and host microtubules were labelled with an anti-α-tubulin antibody, the PVM was labelled with an anti-UIS4 antibody, and nuclei were stained with DAPI. Elongated parasite microtubule structures were observed under this condition. **(G)** Representative maximum-intensity projection of a PS-ExM confocal z-stack of an infected HeLa cell treated with 0.5 µM oryzalin and fixed at 48 hpi, showing elongated LSPMB and hemi-spindle structures. Scale bars in **(F, G)**, 5 µm. **(H)** Quantification of combined LSPMB and hemi-spindle length in DMSO- and 0.5 µM oryzalin-treated parasites. The mean measured length increased from approximately 11 µm in DMSO-treated parasites to approximately 37 µm following oryzalin treatment. Each dot represents a measurement from a single cell. Data are presented as mean ± SD. Statistical analysis was performed using a two-tailed unpaired *t*-test. ns, not significant; *p ≤ 0.05; **p ≤ 0.01; ***p ≤ 0.001.

**Figure 2–figure supplement 2.**
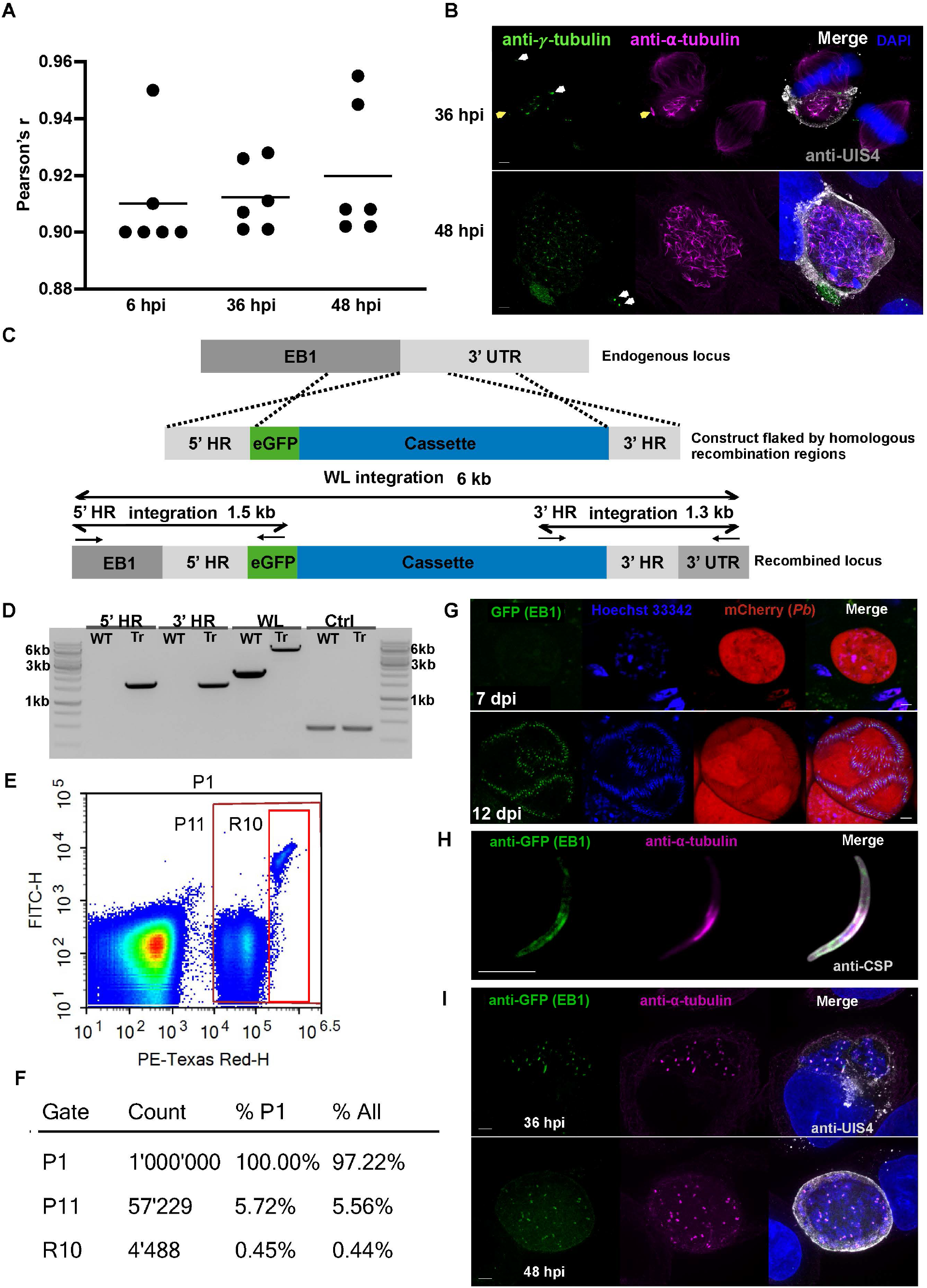
*Plasmodium* end-binding protein 1 is expressed during mosquito and liver stages. **(A)** Pearson correlation coefficients between anti-PolyE and anti-α-tubulin antibody signals associated with the LSPMB at 6, 36 and 48 hpi. The two signals showed a strong positive spatial correlation, with a mean Pearson’s *r* of approximately 0.92 across the analyzed time points. Each dot represents a parasite from a triplicate independent experiment. Data are shown as the mean. The LSPMB region in the parasite was used to calculate Pearson’s r. **(B)** Maximum-intensity projections of PS-ExM confocal z-stacks of infected HeLa cells fixed at 36 and 48 hpi. Samples were labelled with antibodies against γ-tubulin (green), α-tubulin (magenta) and UIS4 (grey); nuclei were stained with DAPI (blue). White arrowheads indicate host-cell centrosomes, and yellow arrowheads indicate anti-γ-tubulin-positive parasite hemi-spindle poles associated with the LSPMB. Scale bars, 10 µm. **(C)** Schematic of the double homologous recombination construct used to tag the C-terminus of *Plasmodium berghei* EB1 (*PBANKA_0405600*) with eGFP. The selection cassette (blue) contains the *P. berghei DHFR* 3′ UTR, the EF1α promoter, human *DHFR*, yeast *FCU*, and the *PBANKA_0405600* 3′ UTR. The construct was flanked by a 5′ homologous region containing the C-terminal portion of the EB1 coding sequence and a 3′ homologous region derived from the EB1 3′ UTR. The construct was transfected into the PbmCherry 1868 parental line. **(D)** Integration-PCR analysis confirming insertion of the eGFP-tagging construct into the EB1 locus. The expected products were 1.5 kb for the 5′ integration PCR, 1.3 kb for the 3′ integration PCR, and 6 kb for the whole locus (WL). Primer sequences are listed in Appendix 1–Table 1. Tr, transgenic line; WT, parental line; Ctrl, internal PCR control. The outer lanes contain a 1 kb DNA ladder. **(E)** Representative flow-cytometry profile of erythrocytes infected with EB1–eGFP parasites. P11 identifies mCherry-positive infected erythrocytes based on PE-Texas Red fluorescence, and R10 identifies the eGFP-positive subset within the mCherry-positive population based on FITC fluorescence. The P1 gate represents the total population of erythrocytes analyzed. **(F)** Numerical summary of the flow-cytometry gating shown in **(E)**, indicating event counts and percentages within the respective parental gates and among all recorded events. **(G)** Representative live-cell confocal images of EB1–eGFP-expressing *P. berghei* oocysts at 7 and 12 days after the infectious blood meal. EB1–eGFP is shown in green, parasite-expressed mCherry in red, and Hoechst 33342-stained nuclei in blue. Scale bars, 5 µm. **(H)** Representative confocal immunofluorescence image of salivary gland sporozoites expressing the EB1-GFP labelled with antibodies against GFP (green), α-tubulin (magenta) and circumsporozoite protein (CSP; grey). Scale bar, 10 µm. **(I)** Maximum-intensity projections of PS-ExM confocal z-stacks of HeLa cells infected with EB1–eGFP-expressing parasites at 36 and 48 hpi. Samples were labelled with antibodies against GFP (green), α-tubulin (magenta) and UIS4 (grey); nuclei were stained with DAPI. The anti-GFP signal was detected in association with parasite microtubule structures during liver stage development. Scale bars, 10 µm. Images are representative of three independent experiments.

**Figure 3–figure supplement 1.**
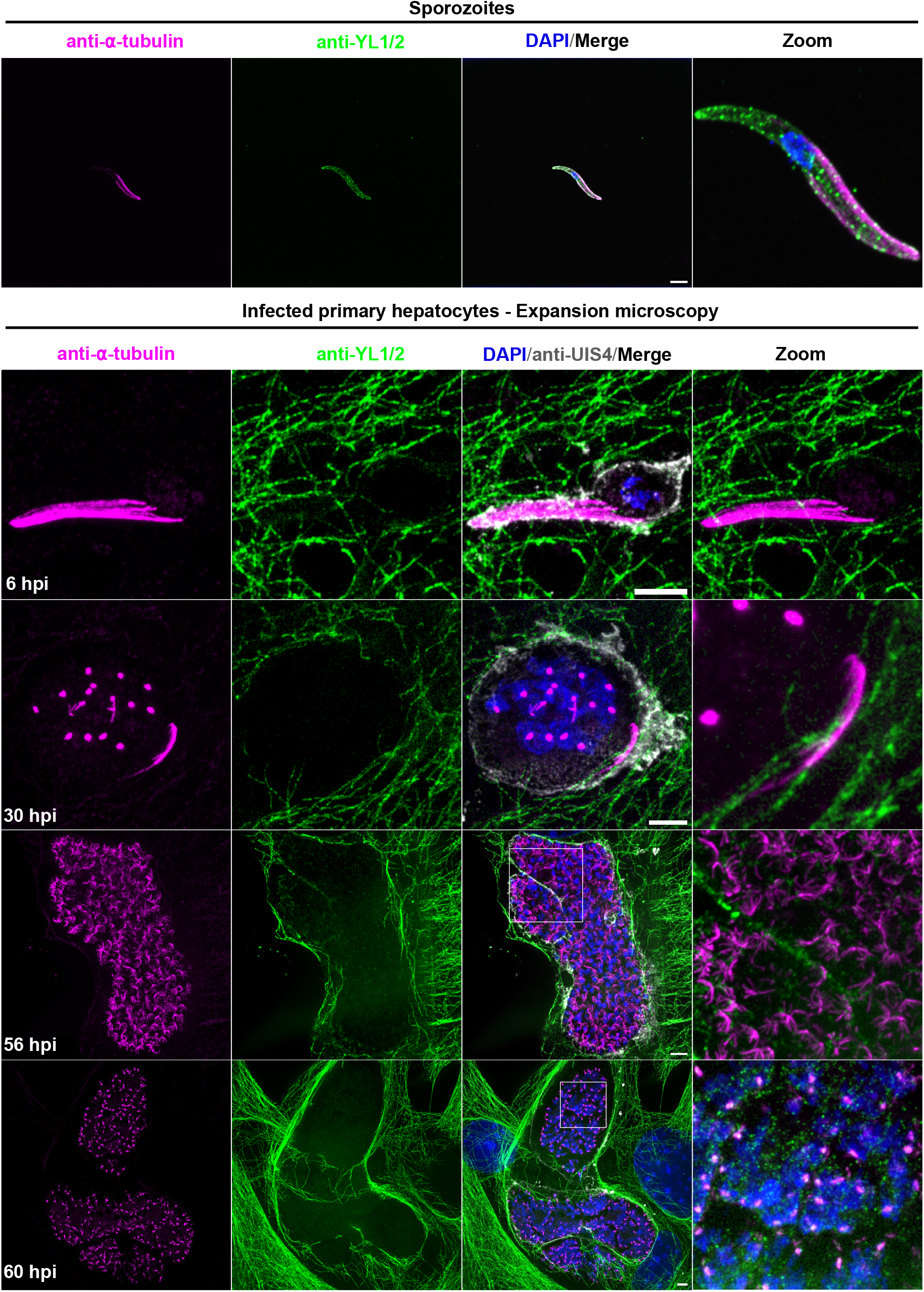
Developmental changes in anti-YL1/2 antibody signals during *Plasmodium* liver stage development. Representative maximum-intensity projections of confocal z-stacks acquired by expansion microscopy from salivary gland sporozoites and infected primary mouse hepatocytes at 6, 30, 56 and 60 hpi. Samples were labelled with antibodies against α-tubulin (magenta) and the C-terminal tyrosine-associated epitope recognized by anti-YL1/2 (green). The PVM was labelled with an anti-UIS4 antibody (grey), and nuclei were stained with DAPI (blue). Merged and magnified views are shown. In extracellular sporozoites, the anti-YL1/2 signal was distributed along the parasite and enriched towards the apical end but showed only limited overlap with the anti-α-tubulin-labelled SSPMTs. This staining pattern differed from the more extensive SSPMT-associated anti-Ty signal detected using antibody T9028 in Figure 3. During early liver stage development, no anti-YL1/2 signal was detected on the anti-α-tubulin-labelled LSPMB above the strong signal associated with host cell microtubules. At 56 and 60 hpi, punctate anti-YL1/2 signals appeared within the parasite. These signals emerged at a similar developmental stage to the punctate anti-Ty signals detected using T9028 but showed a distinct spatial pattern. Scale bars, 10 µm. Images are representative of three independent experiments.

**Figure 4–figure supplement 1.**
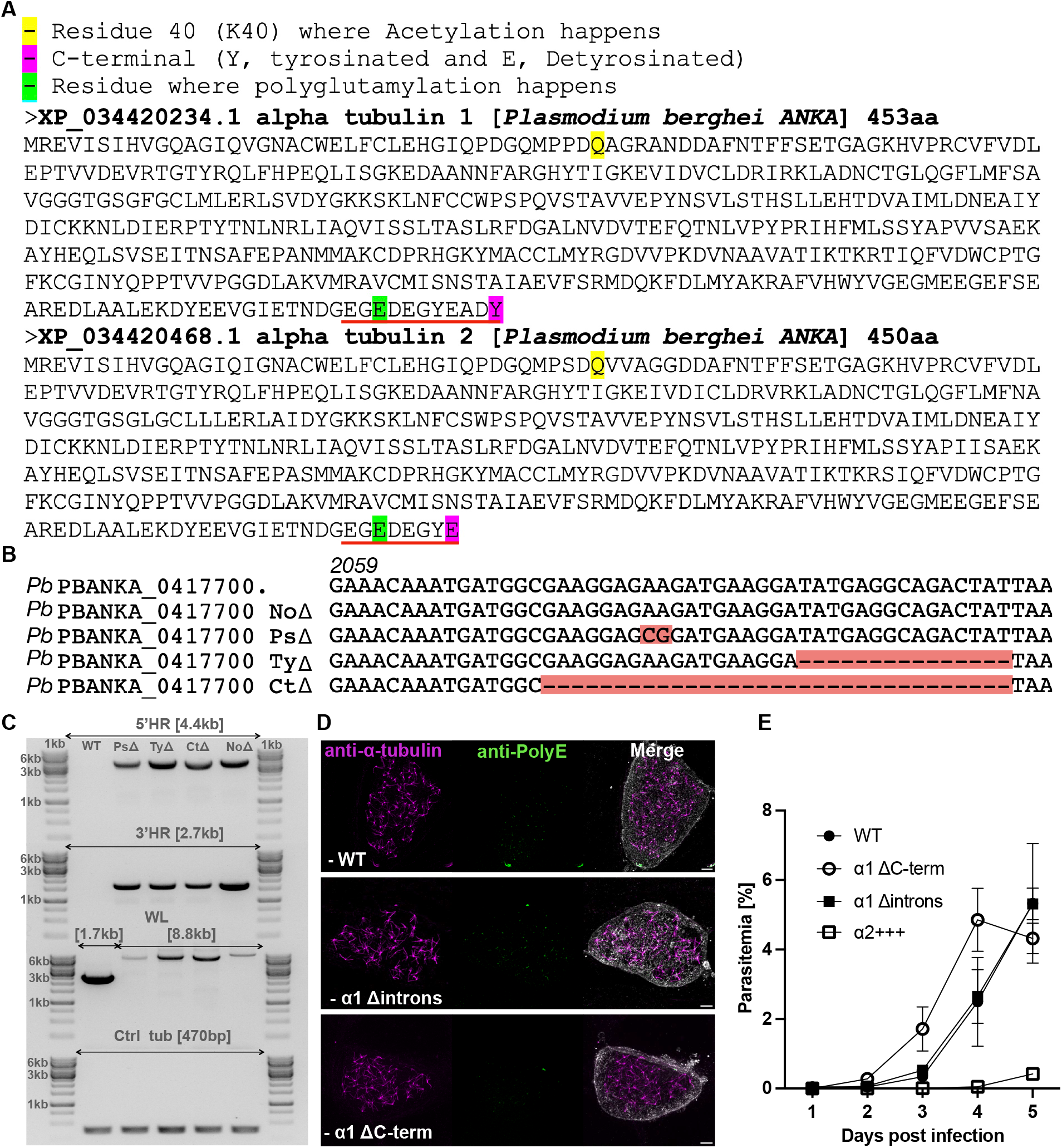
*Plasmodium* α-tubulin proteins and characterization of mutant parasite lines. **(A)** Comparison of *Plasmodium* α1- and α2-tubulin protein sequences, highlighting residues corresponding to canonical tubulin post-translational modification sites. The glutamate at position E445, predicted to be associated with polyglutamylation, is conserved in both isoforms (red). α1-tubulin terminates in tyrosine at Y453, whereas α2-tubulin terminates in glutamate at E450; these residues occupy positions associated with the canonical tyrosination/detyrosination cycle in other α-tubulins (magenta). At the position corresponding to the canonical α-tubulin K40 acetylation site, both *Plasmodium* isoforms contain glutamine and therefore lack the conserved lysine residue (yellow). The position corresponding to the canonical α-tubulin glycylation site is shown in cyan. Human α-tubulin is included for comparison. C-terminal regions associated with tubulin modifications are underlined in red. **(B)** Nucleotide-sequence alignment of *PBANKA_0417700*, showing the mutations introduced into the α1-tubulin mutant parasite lines. Red highlighting indicates substitutions or deletions, and dashes indicate deleted nucleotides. PsΔ contains an E445A substitution at a predicted polyglutamylation-associated residue; TyΔ lacks the C-terminal YEADY motif; CtΔ lacks the complete α1-tubulin C-terminal tail; and NoΔ is a construct control without a coding-sequence mutation. The number 2059 indicates the nucleotide position in *PBANKA_0417700*. **(C)** Integration-PCR analysis of the mutant parasite lines on a 1% agarose gel. The expected products were 4.4 kb for the 5′ integration PCR, 2.7 kb for the 3′ integration PCR, and 8.8 kb for the whole locus (WL) in the recombined parasite lines. The corresponding WT whole-locus product was 1.7 kb. A fragment of the α1-tubulin gene of approximately 470 bp was amplified as a DNA-loading control (Ctrl tub). The outer lanes contain a 1 kb DNA ladder. WT, wild type; PsΔ, E445A substitution; TyΔ, deletion of the C-terminal YEADY motif; CtΔ, deletion of the complete C-terminal tail; NoΔ, construct control without a coding-sequence mutation. **(D)** Maximum-intensity projections of confocal z-stacks of HeLa cells infected with previously generated α1-tubulin mutant parasites, adapted from reference 40. Microtubules were stained with an anti-α-tubulin antibody (magenta), and the anti-PolyE antibody signal is shown in green. Nuclei were stained with DAPI (blue), and the PVM was labelled with an anti-UIS4 antibody (grey). The α1 Δintrons line contains an intronless α1-tubulin gene. The historical α1 ΔC-term line lacks the terminal ADY residues and is distinct from the CtΔ line generated in the present study, in which the complete α1-tubulin C-terminal tail was deleted. Scale bars, 10 µm. **(E)** Parasitemia in mice following intravenous inoculation with 5000 sporozoites of previously generated mutant parasite lines (reference 40). Mice were infected with 5000 sporozoites each. Parasitemia was measured daily from days 1–5 after infection. No statistically significant differences were detected between WT, α1 Δintrons and α1 ΔC-term parasites. In contrast, the α2+++ line, in which α1-tubulin was replaced with α2-tubulin, showed delayed development of detectable blood-stage parasitemia. Three mice per group were analyzed in each of three independent experiments. Data are presented as mean ± SD.

**Figure 5–figure supplement 1.**
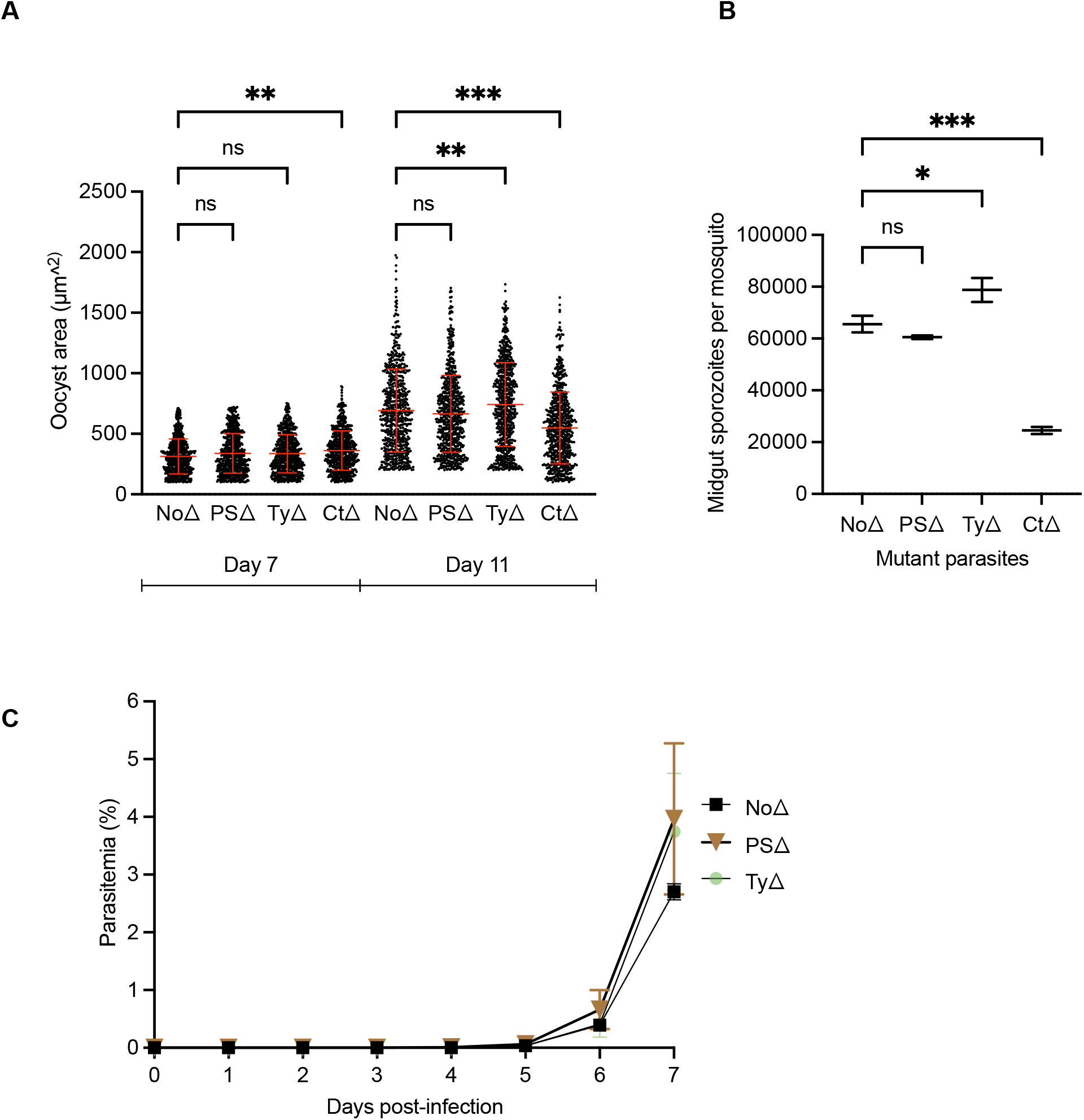
α1-tubulin mutants cand midgut sporozoite production. **(A)** Oocyst areas at days 7 and 11 after the infectious blood meal. At day 7, PsΔ and TyΔ oocyst areas did not differ significantly from NoΔ, whereas CtΔ oocysts were significantly larger than NoΔ oocysts. At day 11, CtΔ oocysts were significantly smaller, whereas TyΔ oocysts were significantly larger, than NoΔ oocysts. Each dot represents one oocyst. Twenty infected mosquitoes per parasite line were dissected in each of three independent experiments. Oocyst areas were analyzed using a mixed-effects model to account for the clustering of multiple oocysts within individual mosquitoes, followed by Dunnett-adjusted comparisons with NoΔ. Red summary bars indicate the median and interquartile range. **(B)** Midgut sporozoite numbers per mosquito at day 14 after the infectious blood meal. Sporozoite numbers did not differ significantly between NoΔ and PsΔ. TyΔ produced significantly more midgut sporozoites, whereas CtΔ produced significantly fewer, than NoΔ. Each point represents the mean number of sporozoites per mosquito from one independent mosquito cohort. Data are presented as mean ± SD from three independent experiments. Statistical analysis was performed using one-way ANOVA followed by Dunnett’s multiple-comparisons test. **(C)** Blood-stage parasitemia following intravenous infection of mice with 5000 salivary gland sporozoites. PsΔ and TyΔ parasites established blood-stage infections with kinetics similar to those of the NoΔ construct control. CtΔ was not included because the very low number of salivary gland sporozoites precluded an equivalent infection experiment. Parasitemia was measured from days 0–7 after infection. Data are presented as mean ± SD for three mice per group from three independent experiments. Statistical comparisons were performed at day 7 using one-way ANOVA followed by Dunnett’s multiple-comparisons test. PsΔ contains an E445A substitution at a predicted polyglutamylation-associated residue; TyΔ lacks the C-terminal YEADY motif associated with the anti-Ty antibody epitope; CtΔ lacks the α1-tubulin C-terminal tail; and NoΔ is a construct control without a coding-sequence mutation. ns, not significant; *p ≤ 0.05; **p ≤ 0.01; ***p ≤ 0.001.

## Supplementary tables

**Appendix 1–Table 1:**
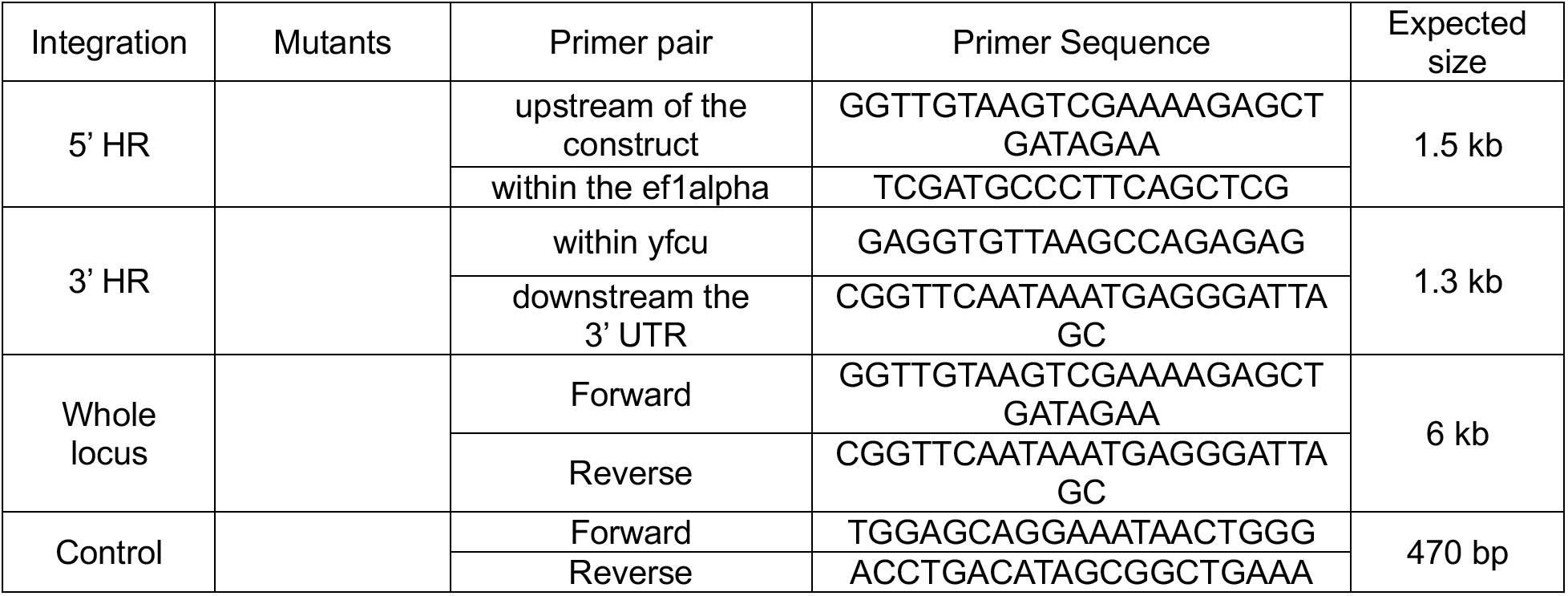
*Plasmodium berghei* EB1egfp parasite integration PCR primers.

**Appendix 1–Table 2:**
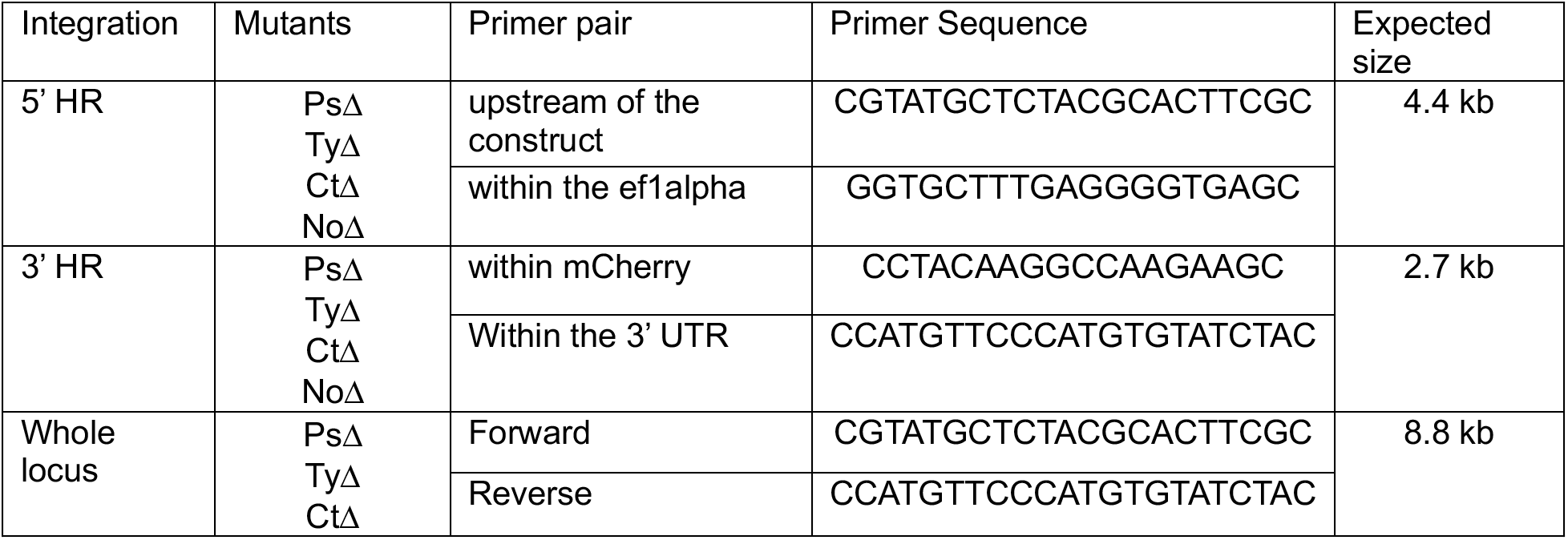
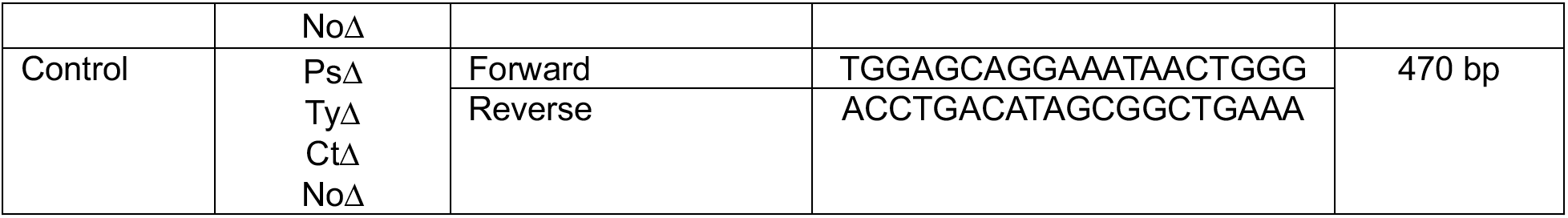
*Plasmodium berghei* α1-tubulin mutant parasite integration PCR primers.

**Appendix 1–Table 3:**
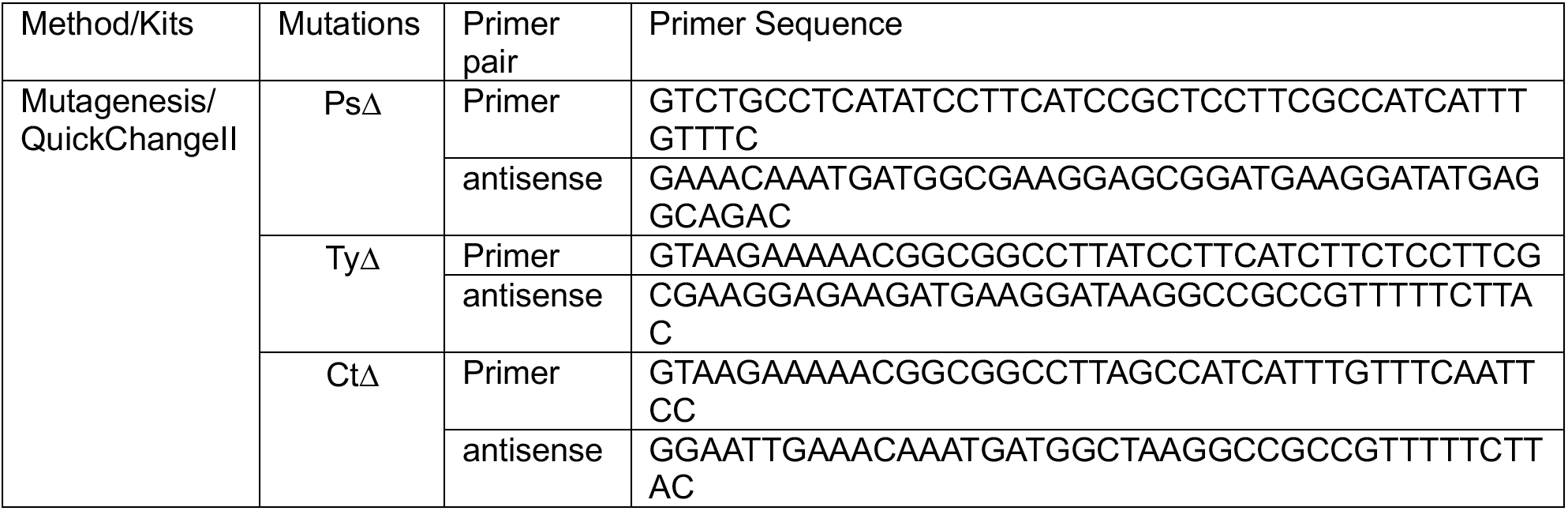
*Plasmodium berghei* α1-tubulin mutagenesis QuickChangeII primer sequences.

### Supplementary videos

**Figure 3–video supplement 1:** Three-dimensional (3D) projection of PS-ExM images showing the LSPMB closely apposed to a hemi-spindle pole in *P. berghei*-infected primary hepatocytes. Fixed and expanded primary hepatocytes were stained with anti-IN105 (anti-PolyE signal, green), anti-α-tubulin (magenta), and anti-UIS4 (PVM, grey). The movie shows the LSPMB is closely associated with the parasite’s hemi-spindle pole. Image analysis was performed using IMARIS 3D and 4D visualization software.

**Figure 4–video supplement 1:** *Plasmodium berghei* salivary gland (SG) sporozoite motility of *P. berghei* NoΔ (control), PsΔ, and TyΔ mutants, as well as hemolymph sporozoites of *P. berghei NoΔ* and CtΔ mutants. *Scale bar: 5 µm*.

## References

1. Wambani J, Okoth P. Impact of Malaria Diagnostic Technologies on the Disease Burden in the Sub-Saharan Africa [Internet]. 2022. doi:10.1155/2022/7324281

2. Poespoprodjo JR, Douglas NM, Ansong D, Kho S, Anstey NM. Malaria. Lancet. 2023 Dec 16;402(10419):2328–45. doi:10.1016/S0140-6736(23)01249-7 PubMed PMID: 37924827.

3. Varo R, Chaccour C, Bassat Q. Update on malaria. Med Clin (Barc). 2020 Nov 13;155(9):395–402. doi:10.1016/J.MEDCLI.2020.05.010 PubMed PMID: 32620355.

4. Venkatesan P. WHO world malaria report 2024. Lancet Microbe. 2025 Apr 1;6(4):101073. doi:10.1016/J.LANMIC.2025.101073 PubMed PMID: 39923782.

5. Tavares J, Formaglio P, Thiberge S, Mordelet E, Van Rooijen N, Medvinsky A, et al. Role of host cell traversal by the malaria sporozoite during liver infection. Journal of Experimental Medicine. 2013 May 6;210(5):905–15. doi:10.1084/JEM.20121130 PubMed PMID: 23610126.

6. Zanghi G, Vaughan AM. Plasmodium vivax pre-erythrocytic stages and the latent hypnozoite. Parasitol Int. 2021 Dec 1;85:102447. doi:10.1016/J.PARINT.2021.102447 PubMed PMID: 34474178.

7. Prudêncio M, Rodriguez A, Mota MM. The silent path to thousands of merozoites: the Plasmodium liver stage. Nature Reviews Microbiology 2006 4:11. 2006 Nov;4(11):849–56. doi:10.1038/nrmicro1529 PubMed PMID: 17041632.

8. Roques M, Bindschedler A, Beyeler R, Heussler VT. Same, same but different: Exploring Plasmodium cell division during liver stage development. PLoS Pathog. 2023 Mar 1;19(3):e1011210. doi:10.1371/JOURNAL.PPAT.1011210 PubMed PMID: 36996035.

9. Shears MJ, Sekhar Nirujogi R, Swearingen KE, Renuse S, Mishra S, Jaipal Reddy P, et al. Proteomic Analysis of Plasmodium Merosomes: The Link between Liver and Blood Stages in Malaria. J Proteome Res. 2019 Sep 6;18(9):3404–18. doi:10.1021/ACS.JPROTEOME.9B00324 PubMed PMID: 31335145.

10. De Niz M, Heussler VT. Rodent malaria models: insights into human disease and parasite biology. Curr Opin Microbiol. 2018 Dec 1;46:93–101. doi:10.1016/J.MIB.2018.09.003 PubMed PMID: 30317152.

11. Sturm A, Amino R, Van De Sand C, Regen T, Retzlaff S, Rennenberg A, et al. Manipulation of host hepatocytes by the malaria parasite for delivery into liver sinusoids. Science (1979). 2006 Sep 1;313(5791):1287–90. doi:10.1126/SCIENCE.1129720 PubMed PMID: 16888102.

12. Mitchison T, Kirschner M. Dynamic instability of microtubule growth. Nature 1984 312:5991. 1984;312(5991):237–42. doi:10.1038/312237a0 PubMed PMID: 6504138.

13. Janke C, Magiera MM. The tubulin code and its role in controlling microtubule properties and functions. Nat Rev Mol Cell Biol. 2020 Jun 1;21(6):307–26. doi:10.1038/S41580-020-0214-3 PubMed PMID: 32107477.

14. Linck R, Fu X, Lin J, Ouch C, Schefter A, Steffen W, et al. Insights into the Structure and Function of Ciliary and Flagellar Doublet Microtubules: TEKTINS, Ca2+-BINDING PROTEINS, AND STABLE PROTOFILAMENTS. J Biol Chem. 2014 Jun 20;289(25):17427. doi:10.1074/JBC.M114.568949 PubMed PMID: 24794867.

15. Owa M, Uchihashi T, Yanagisawa H aki, Yamano T, Iguchi H, Fukuzawa H, et al. Inner lumen proteins stabilize doublet microtubules in cilia and flagella. Nature Communications 2019 10:1. 2019 Mar 8;10(1):1143-. doi:10.1038/s41467-019-09051-x PubMed PMID: 30850601.

16. Nogales E. Structural insights into microtubule function. Annu Rev Biophys Biomol Struct. 2001 Jun 1;30(Volume 30, 2001):397–420. doi:10.1146/ANNUREV.BIOPHYS.30.1.397 PubMed PMID: 11441808.

17. Roll-Mecak A. The Tubulin Code in Microtubule Dynamics and Information Encoding. Dev Cell. 2020 Jul 6;54(1):7–20. doi:10.1016/J.DEVCEL.2020.06.008 PubMed PMID: 32634400.

18. Bodakuntla S, Jijumon AS, Villablanca C, Gonzalez-Billault C, Janke C. Microtubule-Associated Proteins: Structuring the Cytoskeleton. Trends Cell Biol. 2019 Oct 1;29(10):804–19. doi:10.1016/j.tcb.2019.07.004 PubMed PMID: 31416684.

19. Lee G. Non-motor microtubule-associated proteins. Curr Opin Cell Biol. 1993;5(1):88–91. doi:10.1016/S0955-0674(05)80013-4 PubMed PMID: 8448035.

20. Peng N, Nakamura F. Microtubule-associated proteins and enzymes modifying tubulin. Cytoskeleton. 2023 Mar 1;80(3–4):60–76. doi:10.1002/CM.21748 PubMed PMID: 36798013.

21. Fu G, Yan S, Khoo CJ, Chao VC, Liu Z, Mukhi M, et al. Integrated regulation of tubulin tyrosination and microtubule stability by human α-tubulin isotypes. Cell Rep. 2023 Jun 27;42(6):112653. doi:10.1016/J.CELREP.2023.112653 PubMed PMID: 37379209.

22. Gagnon C, White D, Cosson J, Huitorel P, Eddé B, Desbruyères E, et al. The polyglutamylated lateral chain of alpha-tubulin plays a key role in flagellar motility. J Cell Sci. 1996 Jun 1;109(6):1545–53. doi:10.1242/JCS.109.6.1545 PubMed PMID: 8799841.

23. Edde B, Rossier J, Le Caer JP, Prome JC, Desbruyeres E, Gros F, et al. Polyglutamylated alpha-tubulin can enter the tyrosination/detyrosination cycle. Biochemistry. 1992 Feb 1;31(2):403–10. doi:10.1021/BI00117A014 PubMed PMID: 1370628.

24. Ruse CI, Chin HG, Pradhan S. Polyglutamylation: biology and analysis. Amino Acids. 2022 Apr 1;54(4):529–42. doi:10.1007/S00726-022-03146-4 PubMed PMID: 35357568.

25. Zheng P, Obara CJ, Szczesna E, Nixon-Abell J, Mahalingan KK, Roll-Mecak A, et al. ER proteins decipher the tubulin code to regulate organelle distribution. Nature. 2022 Jan 6;601(7891):132–8. doi:10.1038/S41586-021-04204-9 PubMed PMID: 34912111.

26. Mukai M, Ikegami K, Sugiura Y, Takeshita K, Nakagawa A, Setou M. Recombinant mammalian tubulin polyglutamylase TTLL7 performs both initiation and elongation of polyglutamylation on beta-tubulin through a random sequential pathway. Biochemistry. 2009 Feb 10;48(5):1084–93. doi:10.1021/BI802047Y PubMed PMID: 19152315.

27. Janke C, Rogowski K, Wloga D, Regnard C, Kajava A V., Strub JM, et al. Tubulin polyglutamylase enzymes are members of the TTL domain protein family. Science. 2005 Jun 17;308(5729):1758–62. doi:10.1126/SCIENCE.1113010 PubMed PMID: 15890843.

28. Fennell BJ, Al-shatr ZA, Bell A. Isotype expression, post-translational modification and stage-dependent production of tubulins in erythrocytic Plasmodium falciparum. Int J Parasitol. 2008 Apr 1;38(5):527–39. doi:10.1016/J.IJPARA.2007.09.005 PubMed PMID: 17977543.

29. Rawlings DJ, Fujioka H, Fried M, Keister DB, Aikawa M, Kaslow DC. α-Tubulin II is a male-specific protein in Plasmodium falciparum. Mol Biochem Parasitol. 1992 Dec 1;56(2):239–50. doi:10.1016/0166-6851(92)90173-H PubMed PMID: 1484548.

30. Neff NF, Thomas JH, Grisafi P, Botstein D. Isolation of the beta-tubulin gene from yeast and demonstration of its essential function in vivo. Cell. 1983;33(1):211–9. doi:10.1016/0092-8674(83)90350-1 PubMed PMID: 6380751.

31. Baum P, Thorner J, Honig L. Identification of tubulin from the yeast Saccharomyces cerevisiae. Proc Natl Acad Sci U S A. 1978;75(10):4962–6. doi:10.1073/PNAS.75.10.4962 PubMed PMID: 368805.

32. Hirst WG, Fachet D, Kuropka B, Weise C, Saliba KJ, Reber S. Purification of functional Plasmodium falciparum tubulin allows for the identification of parasite-specific microtubule inhibitors. Current Biology. 2022 Feb 28;32(4):919–926.e6. doi:10.1016/J.CUB.2021.12.049 PubMed PMID: 35051355.

33. Holloway SP, Gerousis M, Delves CJ, Sims PFG, Scaife JG, Hyde JE. The tubulin genes of the human malaria parasite Plasmodium falciparum, their chromosomal location and sequence analysis of the α-tubulin II gene. Mol Biochem Parasitol. 1990 Dec 1;43(2):257–70. doi:10.1016/0166-6851(90)90150-K PubMed PMID: 2090947.

34. Fennell BJ, Naughton JA, Dempsey E, Bell A. Cellular and molecular actions of dinitroaniline and phosphorothioamidate herbicides on Plasmodium falciparum: tubulin as a specific antimalarial target. Mol Biochem Parasitol. 2006 Feb;145(2):226–38. doi:10.1016/J.MOLBIOPARA.2005.08.020 PubMed PMID: 16406111.

35. Dow GS, Armson A, Boddy MR, Itenge T, McCarthy D, Parkin JE, et al. Plasmodium: assessment of the antimalarial potential of trifluralin and related compounds using a rat model of malaria, Rattus norvegicus. Exp Parasitol. 2002 Mar 1;100(3):155–60. doi:10.1016/S0014-4894(02)00016-4 PubMed PMID: 12173400.

36. Dempsey E, Prudêncio M, Fennell BJ, Gomes-Santos CS, Barlow JW, Bell A. Antimitotic herbicides bind to an unidentified site on malarial parasite tubulin and block development of liver-stage Plasmodium parasites. Mol Biochem Parasitol. 2013 Apr;188(2):116–27. doi:10.1016/J.MOLBIOPARA.2013.03.001 PubMed PMID: 23523992.

37. Kooij TWA, Franke-Fayard B, Renz J, Kroeze H, Van Dooren MW, Ramesar J, et al. Plasmodium berghei alpha-tubulin II: a role in both male gamete formation and asexual blood stages. Mol Biochem Parasitol. 2005 Nov;144(1):16–26. doi:10.1016/J.MOLBIOPARA.2005.07.003 PubMed PMID: 16115694.

38. Holloway SP, Sims PFG, Delves CJ, Scaife JG, Hyde JE. Isolation of alpha-tubulin genes from the human malaria parasite, Plasmodium falciparum: sequence analysis of alpha-tubulin. Mol Microbiol. 1989;3(11):1501–10. doi:10.1111/J.1365-2958.1989.TB00136.X PubMed PMID: 2693901.

39. Spreng B, Fleckenstein H, Kübler P, Di Biagio C, Benz M, Patra P, et al. Microtubule number and length determine cellular shape and function in Plasmodium. EMBO J. 2019 Aug 24;38(15). 10.15252/embj.2018100984 PubMed PMID: 31368598.

40. Zhang G, Niu G, Hooker–Romera D, Shabani S, Ramelow J, Wang X, et al. Targeting plasmodium α-tubulin-1 to block malaria transmission to mosquitoes. Front Cell Infect Microbiol. 2023;13:1132647. doi:10.3389/FCIMB.2023.1132647 PubMed PMID: 37009496.

41. Valenstein ML, Roll-Mecak A. Graded Control of Microtubule Severing by Tubulin Glutamylation. Cell. 2016 Feb 25;164(5):911–21. doi:10.1016/J.CELL.2016.01.019 PubMed PMID: 26875866.

42. Bonnet C, Boucher D, Lazereg S, Pedrotti B, Islam K, Denoulet P, et al. Differential Binding Regulation of Microtubule-associated Proteins MAP1A, MAP1B, and MAP2 by Tubulin Polyglutamylation. Journal of Biological Chemistry. 2001 Apr 20;276(16):12839–48. doi:10.1074/JBC.M011380200 PubMed PMID: 11278895.

43. Bertiaux E, Balestra AC, Bournonville L, Louvel V, Maco B, Soldati-Favre D, et al. Expansion microscopy provides new insights into the cytoskeleton of malaria parasites including the conservation of a conoid. PLoS Biol. 2021 Mar 11;19(3). doi:10.1371/JOURNAL.PBIO.3001020 PubMed PMID: 33705377.

44. Hentzschel F, Binder AM, Dorner LP, Herzel L, Nuglisch F, Sema M, et al. Microtubule inner proteins of Plasmodium are essential for transmission of malaria parasites. Proceedings of the National Academy of Sciences. 2025 Feb 11;122(6):e2421737122. doi:10.1073/PNAS.2421737122

45. Atchou K, Berger BM, Heussler V, Ochsenreiter T. Pre-gelation staining expansion microscopy for visualisation of the Plasmodium liver stage. J Cell Sci. 2023;136(22):jcs261377. doi:10.1242/JCS.261377/VIDEO-1

46. Liffner B, Absalon S. Expansion microscopy of apicomplexan parasites. Mol Microbiol. 2024 Apr 1;121(4):619–35. doi:10.1111/MMI.15135 PubMed PMID: 37571814.

47. Liffner B, Diaz AKC, Blauwkamp J, Anaguano D, Frolich S, Muralidharan V, et al. Atlas of Plasmodium falciparum intraerythrocytic development using expansion microscopy. Elife. 2023 Dec 18;12. doi:10.7554/ELIFE.88088 PubMed PMID: 38108809.

48. Ferreira JL, Pražák Vojtěch, Vasishtan Daven, Siggel Marc, Hentzschel Franziska, Binder AM, et al. Variable microtubule architecture in the malaria parasite. Nat Commun. 2023 Dec 1;14(1):1216. doi:10.1038/S41467-023-36627-5 PubMed PMID: 36869034.

49. Aguayo-Ortiz R, Dominguez L. Unveiling the Possible Oryzalin-Binding Site in the α-Tubulin of Toxoplasma gondii. ACS Omega. 2022 Jun 7;7(22):18434–42. doi:10.1021/ACSOMEGA.2C00729

50. Beyeler R, Jordan M, Dorner L, He B, Cyrklaff M, Roques M, et al. Putative prefoldin complex subunit 5 of Plasmodium berghei is crucial for microtubule formation and parasite development in the mosquito. Mol Microbiol. 2024 Mar 1;121(3):481–96. doi:10.1111/MMI.15196 PubMed PMID: 38009402.

51. Roques M, Stanway RR, Rea EI, Markus R, Brady D, Holder AA, et al. Plasmodium centrin PbCEN-4 localizes to the putative MTOC and is dispensable for malaria parasite proliferation. Biol Open. 2019 Jan 1;8(1):bio036822. doi:10.1242/BIO.036822/VIDEO-5

52. Zeeshan M, Shilliday F, Liu T, Abel S, Mourier T, Ferguson DJP, et al. Plasmodium kinesin-8X associates with mitotic spindles and is essential for oocyst development during parasite proliferation and transmission. PLoS Pathog. 2019;15(10):e1008048. doi:10.1371/JOURNAL.PPAT.1008048 PubMed PMID: 31600347.

53. Mauer S, Camargo N, Abatiyow BA, Gargaro OR, Kappe SHI, Kumar S. Plasmodium microtubule-binding protein EB1 is critical for partitioning of nuclei in male gametogenesis. mBio. 2023 Aug 31;14(4):e00822–23. doi:10.1128/MBIO.00822-23 PubMed PMID: 37535401.

54. Yang S, Cai M, Huang J, Zhang S, Mo X, Jiang K, et al. EB1 decoration of microtubule lattice facilitates spindle-kinetochore lateral attachment in Plasmodium male gametogenesis. Nature Communications 2023 14:1. 2023 May 19;14(1):1–20. doi:10.1038/s41467-023-38516-3 PubMed PMID: 37208365.

55. Zeeshan M, Rea E, Abel S, Vukušić K, Markus R, Brady D, et al. Plasmodium ARK2 and EB1 drive unconventional spindle dynamics, during chromosome segregation in sexual transmission stages. Nature Communications 2023 14:1. 2023 Sep 13;14(1):1–19. doi:10.1038/s41467-023-41395-3 PubMed PMID: 37704606.

56. Magiera MM, Janke C. Investigating Tubulin Posttranslational Modifications with Specific Antibodies. Methods Cell Biol. 2013 Jan 1;115:247–67. doi:10.1016/B978-0-12-407757-7.00016-5 PubMed PMID: 23973077.

57. Oakley BR. gamma-Tubulin. Curr Top Dev Biol. 2000;49:27–54. doi:10.1016/S0070-2153(99)49003-9 PubMed PMID: 11005013.

58. Maessen S, Wesseling JG, Smits MA, Konings RNH, Schoenmakers JGG. The γ-tubulin gene of the malaria parasite Plasmodium falciparum. Mol Biochem Parasitol. 1993 Jul 1;60(1):27–35. doi:10.1016/0166-6851(93)90025-S PubMed PMID: 8366893.

59. Haase R, Puthenpurackal A, Maco B, Guérin A, Soldati-Favre D. γ-tubulin complex controls the nucleation of tubulin-based structures in Apicomplexa. Mol Biol Cell. 2024 Sep 1;35(9):ar121. doi:10.1091/MBC.E24-03-0100 PubMed PMID: 39046777.

60. Barik S, Andrews J. Host–Parasite Interactions in Toxoplasma gondii-Infected Cells: Roles of Mitochondria, Microtubules, and the Parasitophorous Vacuole. Int J Mol Sci. 2024 Dec 1;25(24):13459. doi:10.3390/IJMS252413459 PubMed PMID: 39769222.

61. Burda PC, Caldelari R, Heussler VT. Manipulation of the Host Cell Membrane during Plasmodium Liver Stage Egress. mBio. 2017 Mar 1;8(2). doi:10.1128/MBIO.00139-17 PubMed PMID: 28400525.

62. Webster DR, Gundersen GG, Bulinski JC, Borisy GG. Differential turnover of tyrosinated and detyrosinated microtubules. Proc Natl Acad Sci U S A. 1987;84(24):9040–4. doi:10.1073/PNAS.84.24.9040 PubMed PMID: 3321065.

63. Arregui C, Busciglio J, Caceres A, Barra HS. Tyrosinated and detyrosinated microtubules in axonal processes of cerebellar macroneurons grown in culture. J Neurosci Res. 1991;28(2):171–81. doi:10.1002/JNR.490280204 PubMed PMID: 1674546.

64. Atchou K, Caldelari R, Roques M, Schmuckli-Maurer J, Beyeler R, Heussler V. Expanding the fluorescent toolkit: Blue fluorescent protein-expressing Plasmodium berghei for enhanced multiplex microscopy. Portugaliza HP, editor. PLoS One. 2025 Mar 3;20(3):e0308055. doi:10.1371/JOURNAL.PONE.0308055 PubMed PMID: 40029851.

65. Hopp CS, Chiou K, Ragheb DRT, Salman AM, Khan SM, Liu AJ, et al. Longitudinal analysis of plasmodium sporozoite motility in the dermis reveals component of blood vessel recognition. Elife. 2015 Aug 13;4(AUGUST2015). doi:10.7554/ELIFE.07789 PubMed PMID: 26271010.

66. Beyer K, Kracht S, Kehrer J, Singer M, Klug D, Frischknecht F. Limited Plasmodium sporozoite gliding motility in the absence of TRAP family adhesins. Malar J. 2021 Dec 1;20(1):1–12. doi:10.1186/S12936-021-03960-3/FIGURES/5 PubMed PMID: 34717635.

67. Klug D, Frischknecht F. Motility precedes egress of malaria parasites from oocysts. Elife. 2017 Jan 24;6. doi:10.7554/ELIFE.19157 PubMed PMID: 28115054.

68. Braumann F, Klug D, Kehrer J, Song G, Feng J, Springer TA, et al. Conformational change of Plasmodium TRAP is essential for sporozoite migration and transmission. EMBO Rep. 2023 Jul 5;24(7). doi:10.15252/EMBR.202357064 PubMed PMID: 37306042.

69. Douglas RG, Reinig M, Neale M, Frischknecht F. Screening for potential prophylactics targeting sporozoite motility through the skin. Malar J. 2018 Aug 31;17(1). doi:10.1186/S12936-018-2469-0 PubMed PMID: 30170589.

70. Meis JFGM, Verhave JP, Jap PHK, Meuwissen JHET. Transformation of sporozoites of Plasmodium berghei into exoerythrocytic forms in the liver of its mammalian host. Cell Tissue Res. 1985 Aug;241(2):353–60. doi:10.1007/BF00217180/METRICS PubMed PMID: 3896506.

71. Short B. Polyglutamylation makes the cut. J Cell Biol. 2010 Jun 14;189(6):920. doi:10.1083/JCB.1896ITI1

72. Genova M, Grycova L, Puttrich V, Magiera MM, Lansky Z, Janke C, et al. Tubulin polyglutamylation differentially regulates microtubule*-*interacting proteins. EMBO J. 2023 Mar;42(5):e112101. doi:10.15252/EMBJ.2022112101 PubMed PMID: 36636822.

73. Pathak NH, Drummond IA. Polyglutamylation and the fleer gene. Methods Cell Biol. 2009;94:317–32. doi:10.1016/s0091-679x(08)94016-4 PubMed PMID: 20362098.

74. Jayabalasingham B, Bano N, Coppens I. Metamorphosis of the malaria parasite in the liver is associated with organelle clearance. Cell Res. 2010 Sep;20(9):1043. doi:10.1038/CR.2010.88 PubMed PMID: 20567259.

75. Graewe S, Stanway RR, Rennenberg A, Heussler VT. Chronicle of a death foretold: Plasmodium liver stage parasites decide on the fate of the host cell. FEMS Microbiol Rev. 2012 Jan 1;36(1):111–30. doi:10.1111/J.1574-6976.2011.00297.X PubMed PMID: 22092244.

76. De Niz M, Caldelari R, Kaiser G, Zuber B, Heo W Do, Heussler VT, et al. Hijacking of the host cell Golgi by Plasmodium berghei liver stage parasites. J Cell Sci. 2021 May 1;134(10):jcs252213. doi:10.1242/JCS.252213/VIDEO-4 PubMed PMID: 34013963.

77. O’neal AJ, Butler LR, Rolandelli A, Gilk SD, Pedra JHF. Lipid hijacking: A unifying theme in vector-borne diseases. Elife. 2020 Oct 1;9:1–31. doi:10.7554/ELIFE.61675 PubMed PMID: 33118933.

78. Fraser M, Curtis B, Phillips P, Yates PA, Lam KS, Netzel O, et al. Harnessing cholesterol uptake of malaria parasites for therapeutic applications. EMBO Mol Med. 2024 Jul 15;16(7):1515–32. doi:10.1038/S44321-024-00087-1 PubMed PMID: 38862600.

79. Schmuckli-Maurer J, Bindschedler AF, Wacker R, Würgler OM, Rehmann R, Lehmberg T, et al. Plasmodium berghei liver stage parasites exploit host GABARAP proteins for TFEB activation. Communications Biology 2024 7:1. 2024 Nov 21;7(1):1–13. doi:10.1038/s42003-024-07242-x

80. Morrissette N, Abbaali I, Ramakrishnan C, Hehl AB. The Tubulin Superfamily in Apicomplexan Parasites. Microorganisms. 2023 Mar 1;11(3). doi:10.3390/MICROORGANISMS11030706 PubMed PMID: 36985278.

81. De Niz M, Stanway RR, Wacker R, Keller D, Heussler VT. An ultrasensitive NanoLuc-based luminescence system for monitoring Plasmodium berghei throughout its life cycle. Malaria Journal 2016 15:1. 2016 Apr 21;15(1):1–24. doi:10.1186/S12936-016-1291-9 PubMed PMID: 27102897.

82. Janse CJ, Ramesar J, Waters AP. High-efficiency transfection and drug selection of genetically transformed blood stages of the rodent malaria parasite Plasmodium berghei. Nature Protocols 2006 1:1. 2006 Jun 29;1(1):346–56. doi:10.1038/nprot.2006.53 PubMed PMID: 17406255.

83. Lin J wen, Annoura T, Sajid M, Chevalley-Maurel S, Ramesar J, Klop O, et al. A novel “Gene Insertion/Marker Out” (GIMO) method for transgene expression and gene complementation in rodent malaria parasites. PLoS One. 2011 Dec 27;6(12). doi:10.1371/journal.pone.0029289 PubMed PMID: 22216235.

84. Philip N, Waters AP. Conditional degradation of plasmodium calcineurin reveals functions in parasite colonization of both host and vector. Cell Host Microbe. 2015 Jul 8;18(1):122–31. doi:10.1016/J.CHOM.2015.05.018 PubMed PMID: 26118994.

85. Patzewitz EM, Guttery DS, Poulin B, Ramakrishnan C, Ferguson DJP, Wall RJ, et al. An Ancient Protein Phosphatase, SHLP1, Is Critical to Microneme Development in Plasmodium Ookinetes and Parasite Transmission. Cell Rep. 2013;3(3):622. doi:10.1016/J.CELREP.2013.01.032 PubMed PMID: 23434509.

86. Salman AM, Mogollon CM, Lin JW, Van Pul FJA, Janse CJ, Khan SM. Generation of transgenic rodent malaria parasites expressing human malaria parasite proteins. Methods in Molecular Biology. 2015;1325:257–86. doi:10.1007/978-1-4939-2815-6_21 PubMed PMID: 26450395.

87. Govindarajalu G, Rizvi Z, Kumar D, Sijwali PS. Lyse-Reseal Erythrocytes for Transfection of Plasmodium falciparum. Sci Rep. 2019 Dec 1;9(1). doi:10.1038/S41598-019-56513-9 PubMed PMID: 31882761.

88. Pino P, Caldelari R, Mukherjee B, Vahokoski J, Klages N, Maco B, et al. A multistage antimalarial targets the plasmepsins IX and X essential for invasion and egress. Science (1979). 2017 Oct 27;358(6362):522–8. doi:10.1126/SCIENCE.AAF8675 PubMed PMID: 29074775.

89. Prado M, Eickel N, De Niz M, Heitmann A, Agop-Nersesian C, Wacker R, et al. Long-term live imaging reveals cytosolic immune responses of host hepatocytes against Plasmodium infection and parasite escape mechanisms. Autophagy. 2015;11(9):1561–79. doi:10.1080/15548627.2015.1067361 PubMed PMID: 26208778.

90. Kaur I, Vasudevan A, Rawal P, Tripathi DM, Ramakrishna S, Kaur S, et al. Primary Hepatocyte Isolation and Cultures: Technical Aspects, Challenges and Advancements. Bioengineering. 2023 Feb 1;10(2):131. doi:10.3390/BIOENGINEERING10020131 PubMed PMID: 36829625.

91. Bindschedler A, Schmuckli-Maurer J, Buchser S, Fischer TD, Wacker R, Davalan T, et al. LC3B labeling of the parasitophorous vacuole membrane of Plasmodium berghei liver stage parasites depends on the V-ATPase and ATG16L1. Mol Microbiol. 2024 Jun 1;121(6):1095–111. doi:10.1111/MMI.15259 PubMed PMID: 38574236.

92. Blight J, Sala KA, Atcheson E, Kramer H, Turabi A El, Real E, et al. Dissection-independent production of Plasmodium sporozoites from whole mosquitoes. Life Sci Alliance. 2021 Jul 1;4(7):e202101094. doi:10.26508/LSA.202101094 PubMed PMID: 34135099.

93. Liffner B, Silva TLA e, Vega-Rodriguez J, Absalon S. Mosquito Tissue Ultrastructure-Expansion Microscopy (MoTissU-ExM) enables ultrastructural and anatomical analysis of malaria parasites and their mosquito. BMC Methods 2024 1:1. 2024 Oct 7;1(1):1–15. doi:10.1186/S44330-024-00013-4

94. Rankin KE, Graewe S, Heussler VT, Stanway RR. Imaging liver-stage malaria parasites. Cell Microbiol. 2010 May;12(5):569–79. doi:10.1111/J.1462-5822.2010.01454.X PubMed PMID: 20180802.

