## Appendix 1 for "Remodeling of sporozoite microtubules during *Plasmodium* liver stage development"

**Appendix 1–Table 1:** *Plasmodium berghei* eB1egfp parasites integration PCR primer

| Integration | Mutants | Primer pair | Primer Sequence | Expected size |
| --- | --- | --- | --- | --- |
| 5’HR |  | upstream of the construct | GGTTGTAAGTCGAAAAGAGCTGATAGAA | 1.5kb |
|  |  | within the ef1alpha | TCGATGCCCTTCAGCTCG |  |
| 3’HR |  | within yfcu | GAGGTGTTAAGCCAGAGAG | 1.3kb |
|  |  | downstream the 3’UTR | CGGTTCAATAAATGAGGGATTAGC |  |
| Whole locus |  | Forward | GGTTGTAAGTCGAAAAGAGCTGATAGAA | 6kb |
|  |  | Reverse | CGGTTCAATAAATGAGGGATTAGC |  |
| Control |  | Forward | TGGAGCAGGAAATAACTGGG | 470bp |
|  |  | Reverse | ACCTGACATAGCGGCTGAAA |  |

**Appendix 1–Table 2:** *Plasmodium berghei* ⍺1-tubulin mutant parasites integration PCR primer

| Integration | Mutants | Primer pair | Primer Sequence | Expected size |
| --- | --- | --- | --- | --- |
| 5’HR | PsΔ  TyΔ  CtΔ  NoΔ | upstream of the construct | CGTATGCTCTACGCACTTCGC | 4.4kb |
|  |  | within the ef1alpha | GGTGCTTTGAGGGGTGAGC |  |
| 3’HR | PsΔ  TyΔ  CtΔ  NoΔ | within mCherry | CCTACAAGGCCAAGAAGC | 2.7kb |
|  |  | Within the 3’UTR | CCATGTTCCCATGTGTATCTAC |  |
| Whole locus | PsΔ  TyΔ  CtΔ  NoΔ | Forward | CGTATGCTCTACGCACTTCGC | 8.8kb |
|  |  | Reverse | CCATGTTCCCATGTGTATCTAC |  |
| Control | PsΔ  TyΔ  CtΔ  NoΔ | Forward | TGGAGCAGGAAATAACTGGG | 470bp |
|  |  | Reverse | ACCTGACATAGCGGCTGAAA |  |

**Appendix 1–Table 3:** *Plasmodium berghei* ⍺1-tubulin mutagenesis QuickChangeII primers sequences

| Method/Kits | Mutations | Primer pair | Primer Sequence |
| --- | --- | --- | --- |
| Mutagenesis/  QuickChangeII | PsΔ | Primer | GTCTGCCTCATATCCTTCATCCGCTCCTTCGCCATCATTTGTTTC |
|  |  | antisense | GAAACAAATGATGGCGAAGGAGCGGATGAAGGATATGAGGCAGAC |
|  | TyΔ | Primer | GTAAGAAAAACGGCGGCCTTATCCTTCATCTTCTCCTTCG |
|  |  | antisense | CGAAGGAGAAGATGAAGGATAAGGCCGCCGTTTTTCTTAC |
|  | CtΔ | Primer | GTAAGAAAAACGGCGGCCTTAGCCATCATTTGTTTCAATTCC |
|  |  | antisense | GGAATTGAAACAAATGATGGCTAAGGCCGCCGTTTTTCTTAC |
